# Relaxin/insulin-like family peptide receptor 4 (Rxfp4) expressing hypothalamic neurons modulate food intake and preference in mice

**DOI:** 10.1101/2021.06.26.450020

**Authors:** Jo E Lewis, Orla RM Woodward, Danaé Nuzzaci, Christopher A Smith, Alice E Adriaenssens, Lawrence Billing, Cheryl Brighton, Benjamin U Phillips, John A Tadross, Sarah J Kinston, Ernesto Ciabatti, Berthold Göttgens, Marco Tripodi, David Hornigold, David Baker, Fiona M Gribble, Frank Reimann

## Abstract

Relaxin/insulin-like-family peptide receptor-4 (RXFP4), the cognate receptor for insulin-like peptide 5 (INSL5), has been implicated in feeding behaviour as *Rxfp4* knockout mice display shorter meal durations and reduced high fat diet (HFD) intake. Here, we generated transgenic *Rxfp4*-Cre mice to explore *Rxfp4* expression and physiology. Using this model, we identified *Rxfp4* expression in the central nervous system, including in the ventromedial hypothalamus (VMH). Intra-VMH infusion of INSL5 increased HFD and highly palatable liquid meal intake (HPM) of *ad libitum* fed wildtype mice. Single-cell RNA-sequencing of VMH *Rxfp4*-expressing cells (RXFP4^VMH^) defined a cluster of *Rxfp4*-labelled neurons expressing *Esr1, Tac1* and *Oxtr,* alongside known appetite-modulating neuropeptide receptors (*Mc4r*, *Cckar* and *Nmur2*). Viral tracing demonstrated RXFP4^VMH^ neural projections to the bed nucleus of the stria terminalis, paraventricular hypothalamus, paraventricular thalamus and central nucleus of the amygdala. Utilising designer receptors exclusively activated by designer drugs (DREADDs), we found that whole body chemogenetic inhibition (Di) of *Rxfp4*-expressing cells, mimicking native INSL5-RXFP4 signalling, increased intake of HFD and HPM, whilst activation (Dq), either at whole body level or specifically within the VMH, reduced HFD and HPM intake and altered food preference. Ablating VMH *Rxfp4*-expressing cells recapitulated the lower HFD intake phenotype of *Rxfp4* knockout mice, resulting in reduced body weight. These findings identify a discrete *Rxfp4*-expressing neuronal population as a key regulator of food intake and preference and reveal hypothalamic RXFP4 signalling as a target for feeding behaviour manipulation.

## Introduction

Relaxin/insulin-like-family peptide receptor-4 (RXFP4) is the cognate receptor for insulin-like peptide 5 (INSL5), a member of the relaxin/insulin-like peptide family(1–3). We previously reported an orexigenic effect of exogenously applied INSL5 in mice(4). More recently, we observed a transient increase in food intake when *Insl5* expressing cells were chemogenetically activated (5). This was only revealed when the dominant anorexigenic action of PYY, co-secreted from the same cell population in the distal large intestine (6), was blocked with a NPY2R antagonist (5). Reports that *Insl5* knockout (*Insl5*^-/-^) mice do not display an observable feeding phenotype and that, in some studies, pharmacological administration of INSL5 (both native and PEGylated forms) failed to increase food intake in lean and obese mice (7, 8), have shed some doubt on whether INSL5 plays a physiologically relevant role in the control of food intake. However, *Rxfp4* knockout (*Rxfp4*^-/-^) mice exhibit shorter meal durations, particularly when fed a high fat diet (HFD), and lack the normal preference for HFD over standard chow diet observed in wildtype mice (4). We therefore consider RXFP4 to be a potential target receptor for the manipulation of feeding behaviour.

RXFP4 is a G_αi/o_-coupled receptor identified in 2003 through its homology to relaxin/insulin-like peptide receptor 3, RXFP3 (9, 10). Binding of INSL5 activates downstream signalling pathways including phosphorylation ERK1/2, Akt, p38MAPK and S6RP and reduces cytosolic cAMP levels through inhibition of adenylate cyclase (2). In addition to the selective ligand INSL5, Relaxin-3 can also activate RXFP4, stimulating comparable signalling pathways (2, 11). *Rxfp4* mRNA expression has been reported in the colon, kidney, heart, liver, testes and ovary of both mice and humans (1, 10, 11). To identify and manipulate *Rxfp4*-expressing cells, we developed a new transgenic mouse model in which Cre-recombinase expression is driven by the *Rxfp4* promoter (*Rxfp4*-Cre). Utilising this model, we found clear Cre-dependent reporter expression within the CNS, including the ventromedial hypothalamus (VMH). This contrasts with our previous study, in which we failed to detect *Rxfp4* expression in the central nervous system by qPCR, and in which intracerebroventricular administration of INSL5 failed to increase food intake significantly (4). Revisiting the potential central role of RXFP4 in food intake regulation, we observed that infusion of INSL5 directly into the VMH induced a significant orexigenic response when mice were offered a HFD or a highly palatable liquid Ensure test meal (HPM), but not when offered a standard chow diet. We therefore used the *Rxfp4*-Cre mouse model to explore the role of *Rxfp4*-expressing cells in shaping food intake and preference.

## Results

To investigate a possible role of *Rxfp4*-expressing cells in feeding control, we generated a new BAC transgenic mouse (*Rxfp4*-Cre) model in which Cre-recombinase is expressed under the control of the *Rxfp4*-promoter (Fig. 1a). By crossing *Rxfp4*-Cre mice with fluorescent protein reporter mice (e.g. Rosa26 fxSTOPfx-EYFP (RXFP4^EYFP^)) (Fig. 1b) we observed *Rxfp4* dependent expression in the colon (Fig. 1c), consistent with previous reports, but also detected reporter expression in the central nervous system (CNS) (Suppl. Fig. 1) by GFP immunohistochemistry. Within the CNS we observed reporter expression in the ventromedial hypothalamus (VMH), the accessory olfactory bulb, septofimbrial nucleus, retrosplenial cortex, and the mammillary body medial and lateral parts, as well as in a lower density of cells in the substantia innominata, cortical and central amygdala, periaqueductal grey, and spinal trigeminal tract of the hindbrain (Suppl. Fig 1). Closer examination of the VMH using RXFP4^GCaMP3^ mice revealed transgene expression in the central and ventrolateral parts (VMHc and VMHvl, respectively) extending into the adjacent tuberal nucleus (TN) (Fig. 1d,e), despite our previously reported failure to amplify *Rxfp4* from hypothalamic cDNA by RT-PCR (4). The VMH is a central hub for the regulation of energy balance and integration of diverse nutritionally regulated hormonal and synaptic inputs, and the adjacent TN has also been implicated in feeding behaviour (12, 13). In the VMH, co-staining with NeuN demonstrated GFP expression in mature *Rxfp4* positive neurons (Fig. 1f). Active transcription of *Rxfp4* mRNA in the adult mouse VMH was confirmed by RNAscope (Fig. 1g,h). To test the functional activity of the RXFP4-INSL5 pathway in the *Rxfp4*-expressing cells in the VMH (RXFP4^VMH^), we performed real time monitoring of cAMP levels in RXFP4^VMH^ using ex vivo CAMPER (14) imaging in hypothalamic slices from Rxfp4-Cre x CAMPER (RXFP4^CAMPER^) mice. We found that 14 out of 54 YFP-positive cells showed a clear reduction in the ΔFRET ratio (YFP/CFP), reflecting a reduction in cAMP levels, in response to INSL5 (Fig. 1i), whilst the remaining cells displayed no obvious change in FRET signal in response to INSL5 addition. To test the functional significance of the RXFP4^VMH^ cells in vivo, we treated wildtype ad libitum (AL) fed lean mice with an intra-VMH infusion of INSL5 after a 2h fast. During the light phase, INSL5 had no effect on standard (std) chow intake (Fig. 1j) but in mice which were habituated to the appearance of a HFD or HPM for 1 hour, treatment with INSL5 significantly increased HFD and HPM intake compared to the vehicle control treatment (Fig. 1k,l).

**Figure 1:**
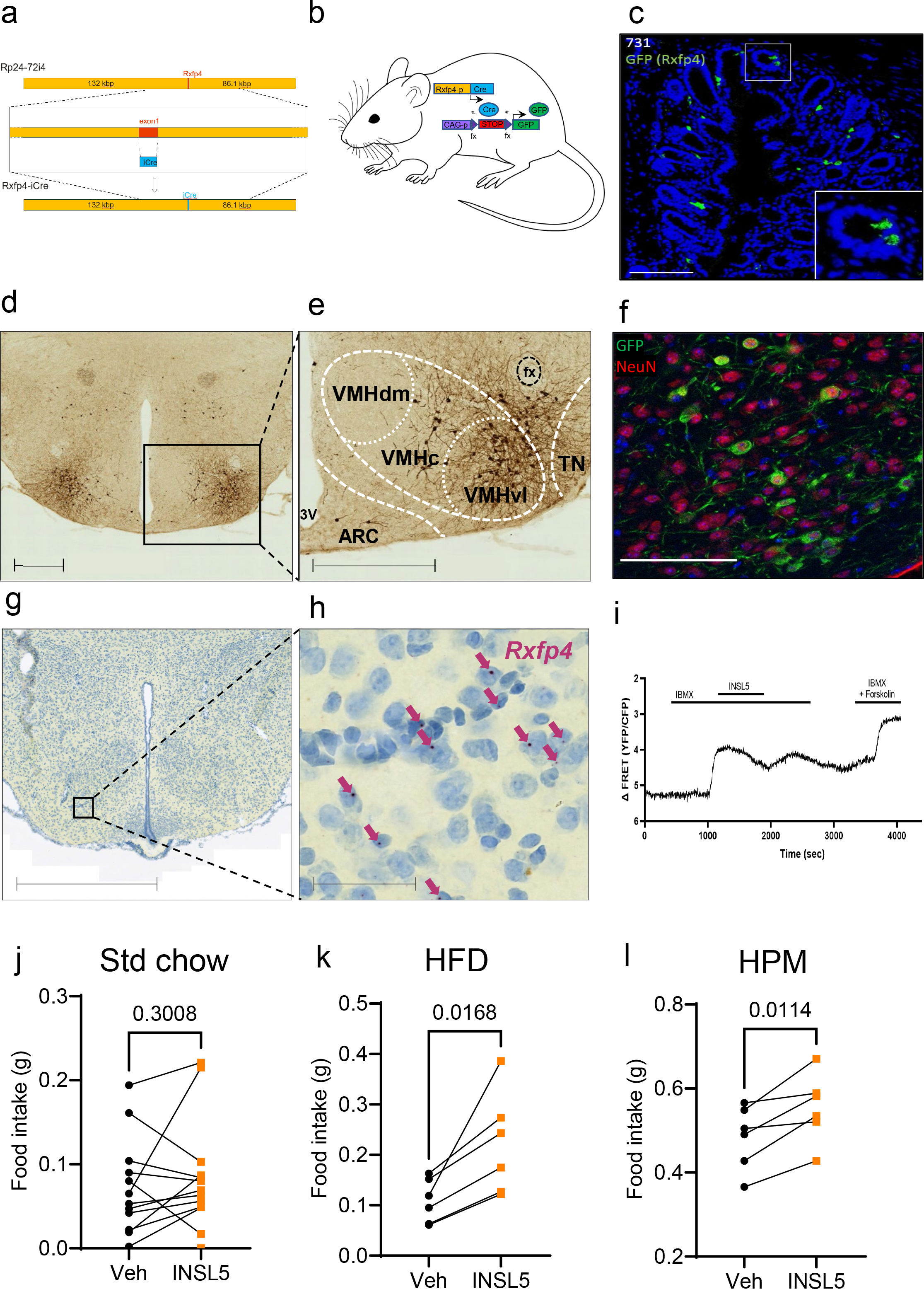
RXFP4 is expressed in the central nervous system. a) Scaled schematic of the bacterial artificial chromosome used to make *Rxfp4*-Cre mice. b) Crossing of *Rxfp4*-Cre mice with GFP-based reporter mice (EYFP or GCaMP3) used to detect *Rxfp4* expression in Fig 1 through GFP immunohistochemistry. c) Representative section from the colon of RXFP4^EYFP^ mice demonstrating *Rxfp4* expression in epithelial cells. Scale bar = 100 μm. d,e) Coronal section of RXFP4^GCaMP3^ mice showing distinct *Rxfp4*-expressing cell clusters in the ventrolateral part of the ventromedial hypothalamus (VMHvl) and adjacent central VMH (VMHc) tuberal nucleus (TN). Scale bars = 500 μm. f) Co-staining for DAPI (blue), GFP (green) and NeuN (red) in the VMH of RXFP4^GCaMP3^ mice. Scale bar = 100 μm. g,h) Coronal sections of C57Bl6 mice labelled for *Rxfp4* mRNA in the VMH using RNAscope. Scale bar = 1 mm and 50 μm in the enlarged image (h). i)Representative trace of a recorded cell in a RXFP4^CAMPER^ slice responding to IBMX (100uM) in the absence and presence of INSL5 (100nM) and Forskolin (10uM), as indicated. 14 out of 54 cells responded to INSL5 with the same pattern (n=5 mice). j, k, l) Food intake of wildtype mice following intra-VMH infusion of vehicle/INSL5 (5 μg) (standard chow t=1.081, p = 0.3008 n=13, HFD (45%) t=3.529 n=6, p = 0.0168, liquid Ensure (HPM) t = 3.902, p = 0.0114 n=6, paired two-tailed t test, animals adapted to the appearance of test meal over the course of two weeks prior to surgery and recovery, testing conducted between 12:00-15:00).

As intra-VMH INSL5 significantly modulated feeding behaviour, we further characterised the *Rxfp4*-expressing cells in this region. Initially, we generated a single cell resolution transcriptomic profile of *Rxfp4*-expressing cells in the hypothalamus. Fluorescent cell populations from the hypothalami of twelve RXFP4^EYFP^ mice were purified by FACS and their transcriptomes analysed by scRNA-Seq. After data normalisation, 350 cells passed quality control filtering. Graph-based clustering analysis revealed that the filtered cells separated into five populations (Fig. 2a). Cluster identities were assigned based on the expression patterns of canonical cell-type markers, identifying microglia (*Tmem119*, *Siglech*, *P2ry12*), neuronal cells (*Snap25, Tubb3, Elavl2*) and endothelial cells (*Dcn*, *Hspg2*), together with smaller clusters of macrophages (*Mrc1*, *Mgl2*) and ependymocytes (*Ccdc153*, *Hdc*) (Fig. 2b). As IHC had suggested neuronal *Rxfp4* expression (Fig1f) and hypothalamic neurons are known to modulate feeding behaviour, we analysed the neuronal cluster in more detail, identifying seven subclusters (Fig. 2c). *Rxfp4*-positive neurons expressed markers for both GABAergic (*Slc32a1*) and glutamatergic (*Slc17a6*) cells (Fig. 2d). Cluster 1 was enriched in markers previously associated with an estrogen receptor (*Esr1*)-positive VMHvl neuronal population (15, 16), including preprotachykinin-1 (*Tac-1*), oxytocin receptor (*Oxtr*), cholecystokinin receptor A (*Cckar*), melanocortin 4 receptor (*Mc4r*) and neuromedin U receptor 2 (*Nmur2*) (Fig. 2d), suggesting crosstalk with known food regulatory networks. Receptors for other established feeding-neuromodulators, like glucagon-like peptide-1 receptor (*Glp1r*) and cholecystokinin receptor B (*Cckbr*), were preferentially expressed in cluster 6 (Fig. 2d).

**Figure 2:**
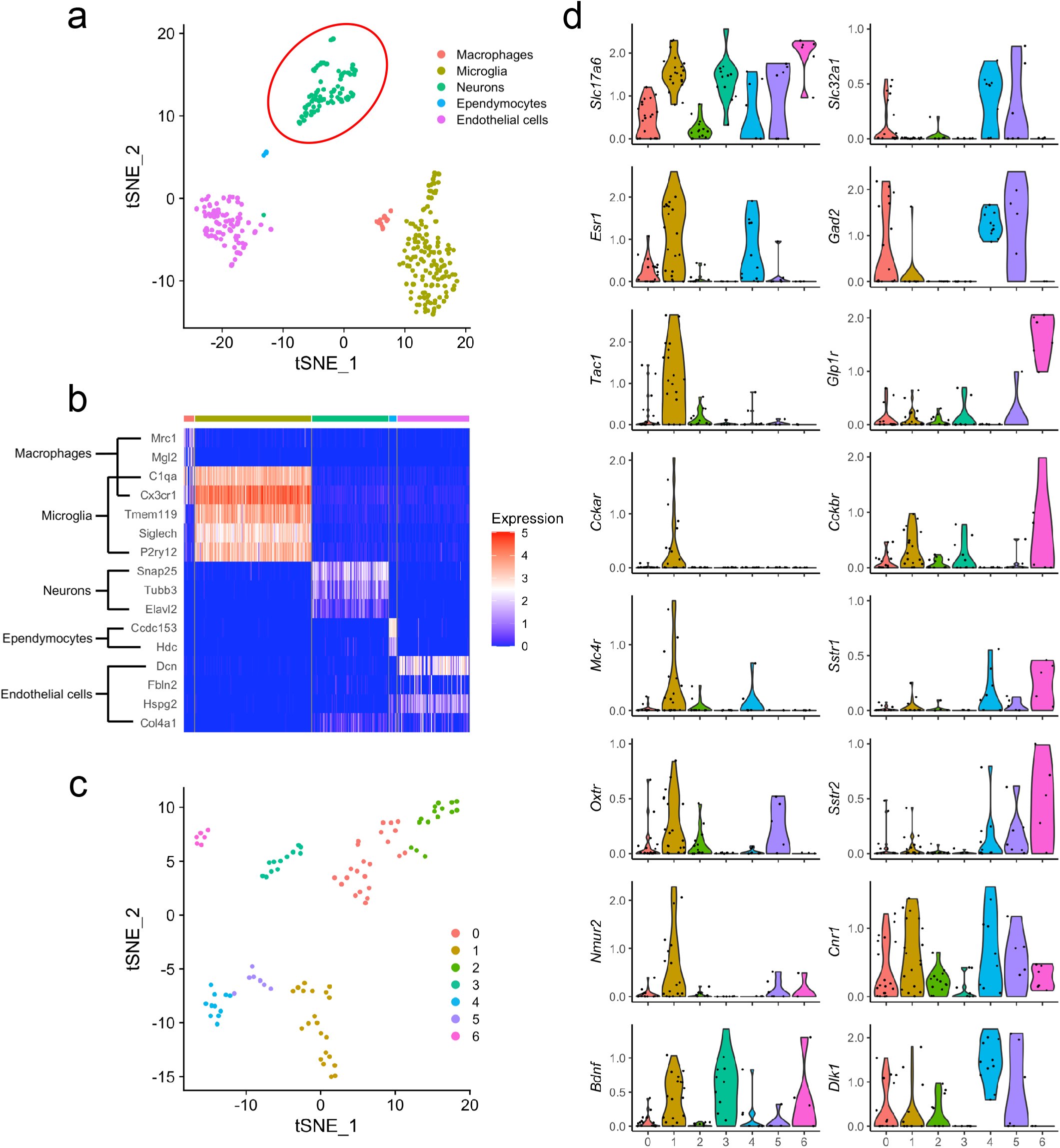
Transcriptomic profiling of hypothalamic *Rxfp4*-expressing cells by scRNAseq. a,b) tSNE visualisation of 350 hypothalamic *Rxfp4*-expressing cells indicates five clusters (a). Cell types were assigned according to the expression of a combination of canonical cell-type markers genes (b). The red circle indicates the neuronal cluster used in further analysis. c) tSNE visualisation of the 95 neuronal cells revealed 7 sub-clusters. D) Violin plots showing expression of marker genes associated with multiple neuronal cell types. All gene expression counts are log-normalised with scale-factor = 10^4^.

We subsequently aimed to establish the neuronal circuitry around RXFP4^VMH^ cells. Anterograde projections were mapped by stereotactically injecting Cre-dependent rAAV8-ChR2-mCherry into the VMH of RXFP4^GCaMP3^mice (17) (Fig.3a,b). Axonal transport of the ChR2-mCherry fusion protein revealed RXFP4^VMH^ projections to multiple regions including the bed nucleus of the stria terminalis (BNST), preoptic area (POA), anteroventral periventricular nucleus (AVPV), arcuate nucleus (ARC), paraventricular hypothalamus (PVH), central nucleus of the amygdala (CeA), periaqueductal grey (PAG, dorsomedial and lateral) and parabrachial nucleus (PBN, lateral and medial) (Fig. 3c, Suppl. Fig. 3). Retrograde projections were assessed after AVV2-TVAeGFP-oG injection followed by Rab-ΔG-EnvA-mCherry (18) injection 21 days later into the VMH of *Rxfp4*-Cre mice (Fig. 3d). The retrograde monosynaptic transport of Rab-mCherry labelled inputs from several nuclei established in feeding regulation, including the ARC, DMH and PVH (Fig. 3d-f). The neuronal circuitry surrounding RXFP4^VMH^ cells is summarised in Fig. 3g.

**Figure 3:**
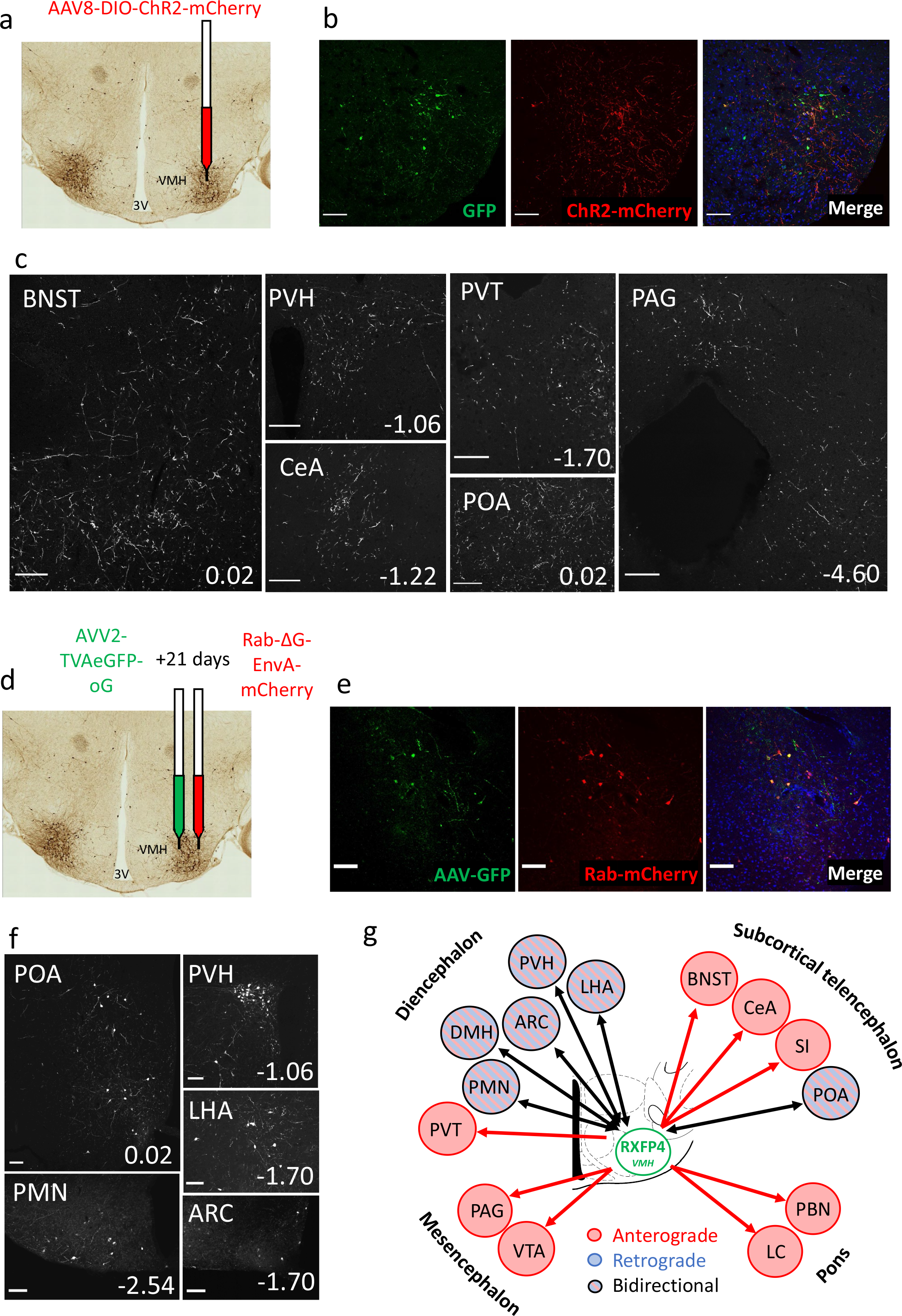
Circuit mapping of hypothalamic *Rxfp4*-expressing cells. a) Schematic illustrating unilateral microinjection of rAAV8-DIO-hChR2(H134R)-mCherry into the VMH of RXFP4^GCaMP3^ mice. B) Immunofluorescence images demonstrating co-localisation of GFP (green) and ChR2-mCherry (red) starter cells in the target region at A/P -1.7mm from bregma. C) Representative images showing ChR2-mCherry-immunoreactive axon terminals in various brain regions (n=3). For each image, distance from bregma (in mm) is indicated at the bottom right. Scale bars = 100 µm. 40x magnification. D) Schematic illustrating unilateral microinjection of AVV2-TVAeGFP-oG into the VMH of *Rxfp4*-Cre mice followed by a unilateral microinjection of Rab-ΔG-EnvA-mCherry 21 days later. E) Immunofluorescence images demonstrating the colocalisation of GFP (green) and Rab-mCherry (red) starter cells in the target region A/P -1.7mm from bregma. F) Representative immunofluorescence images showing Rab-mCherry-immunoreactive cell bodies in various brain regions (n=3). For each image distance from bregma (in mm) is indicated at the bottom right. 20x magnification. G) Schematic illustrating the regions positive for anterograde projections (red arrows), retrograde projections (blue arrows) or bilateral projections (black arrows) from RXFP4^VMH^ cells. Abbreviations: ARC: arcuate nucleus; BNST: bed nucleus of the stria terminalis; CeA: central amygdala; LHA: lateral hypothalamic area; PAG: periaqueductal grey; PMN: premammillary nucleus; POA: Preoptic area; PVH: paraventricular hypothalamus; PVT: paraventricular thalamic nucleus; VMH: ventromedial hypothalamus.

Due to the previously reported altered feeding patterns and macronutrient preferences of *Rxfp4* knock-out mice (4), we further explored the physiology of the RXFP4^VMH^ population by ablating *Rxfp4*-expressing cells using rAAV-DTA injected into the VMH of RXFP4^GCaMP3^ mice (Fig. 4a). Postmortem histological analysis revealed a >90% reduction in RXFP4^VMH^ cells (identified by GFP immunohistochemistry) across the VMH. In the 8 weeks following the surgery, the RXFP4^VMHKO^ mice gained less body weight compared to the control rAAV-mCherry injected mice (Fig. 4c), when fed with a choice of standard chow and HFD in parallel. Five weeks post-surgery the mice were studied in metabolic cages. RXFP4^VMHKO^ mice had a lower respiratory exchange ratio (RER, Fig. 4d), whilst energy expenditure and ambulatory activity (Fig. 4e,f) were unaffected by ablation. Average 24hr food intake was significantly reduced, a consequence of reduced HFD intake (Fig. 4g). Meal duration was unaffected by treatment, however the interval between meals (Fig. 4i) was significantly increased. These data demonstrate the importance of this neuronal population in governing long-term feeding behaviour.

**Figure 4:**
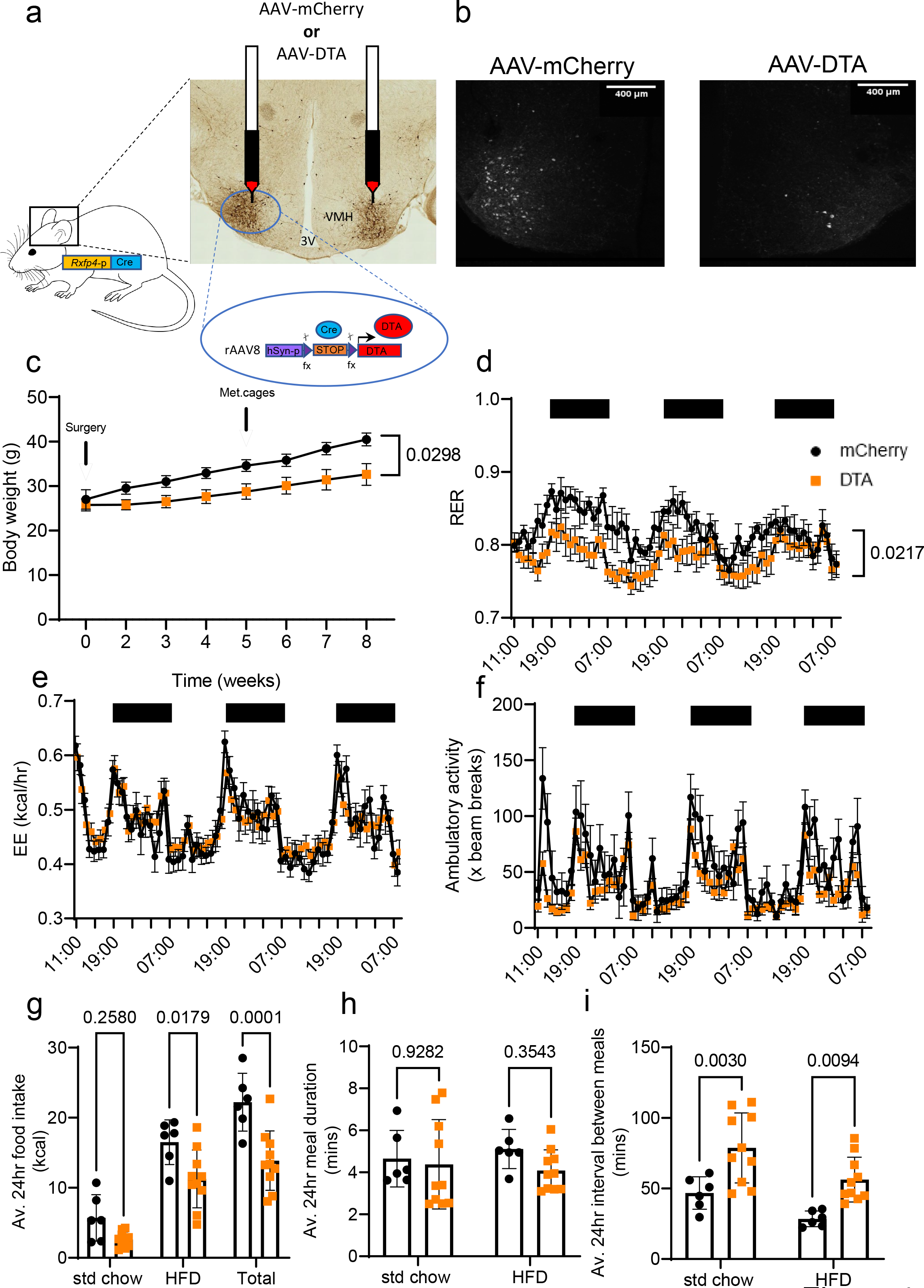
VMH ablation of RXFP4 expressing cells regulates food intake and body weight. a) rAAV-mCherry (control) or rAAV-DTA were bilaterally injected into the ventromedial hypothalamus of *Rxfp4*^GCaMP3^ mice. B) Targeting efficiency was confirmed post perfusion fixation by GFP-immunohistochemistry. Representative images showing efficient ablation in the VMH. C) Body weight following exposure to standard chow and HFD in parallel (effect of time [F(1.423, 22.76) = 44.45, p < 0.0001], effect of treatment [F(1,16) = 5.687, p = 0.03) (n = 8-10 per group). D) RER effect of time [F(6.263, 87.69) = 3.368, p = 0.0044)], effect of treatment [F(1, 14) = 6.669, p = 0.02]. E) Energy expenditure effect of time [F(9.339,130.7) = 20.34, p < 0.0001], effect of treatment [F(1,14) = 0.004, p = 0.95] and f) ambulatory activity effect of time [F(7.723, 100.4) = 11.69, p < 0.0001], effect of treatment [F(1,16) = 3.270, p = 0.09] of mice housed in metabolic cages 5 weeks post-surgery. G) Average 24hr food intake (effect of diet [F(2,42) = 61.97, p < 0.0001) and effect of treatment [F(1.42) = 28.27, p < 0.0001) h) Average 24hr meal duration (effect of diet [F(1,28) = 0.02361, p = 0.88) and effect of treatment [F(1,28) = 1.393, p = 0.25) and i) Average 24hr interval between meals (effect of diet [F(1,28) = 10.08, p = 0.0046) and effect of treatment [F(1,28) = 21.71, p < 0.0001) of mice treated with rAAV-mCherry or rAAV-DTA of RXFP4^VMHKO^ mice in metabolic chambers, when mice had access to a choice between std chow and a HFD (rAAV-mCherry n = 6, rAAV-DTA n =10 respectively, group mean ± SEM).

Given that RXFP4^VHMKO^ mice ate less HFD, yet central Insl5 infusion, which we would predict to inhibit RXFP4-expressing neurons, increased HFD intake, we investigated the effects of acute activation or inhibition of *Rxfp4*-expressing cells. Initially we used a whole-body hM4Di DREADD Cre-reporter (RXFP4^wb-Di^) to mimic the established RXFP4-signalling via pertussis-toxin sensitive Gi pathways (2) (Fig. 5a). During the light phase, activation of Di in *Rxfp4*-expressing cells using CNO had no measurable effect on food intake in AL fed mice(Fig. 5b). However, when animals were habituated to the appearance of a HFD or HPM for 1 hour during the light phase, CNO application resulted in increased food intake (Fig. 5c,d). These results were consistent with the response to infusion of INSL5 into the VMH. To investigate this further, we gave mice housed in metabolic cages the choice between standard chow and HFD. CNO injection during the light phase significantly increased HFD but not chow intake in RXFP4^wb-Di^ mice (Fig. 5e). This effect was transient (Fig. 5f), consistent with the pharmacokinetics of CNO (19), and occurred without any significant differences in ambulatory activity or energy expenditure compared to the vehicle cross-over control (Suppl. Fig. 4a,b).

**Figure 5:**
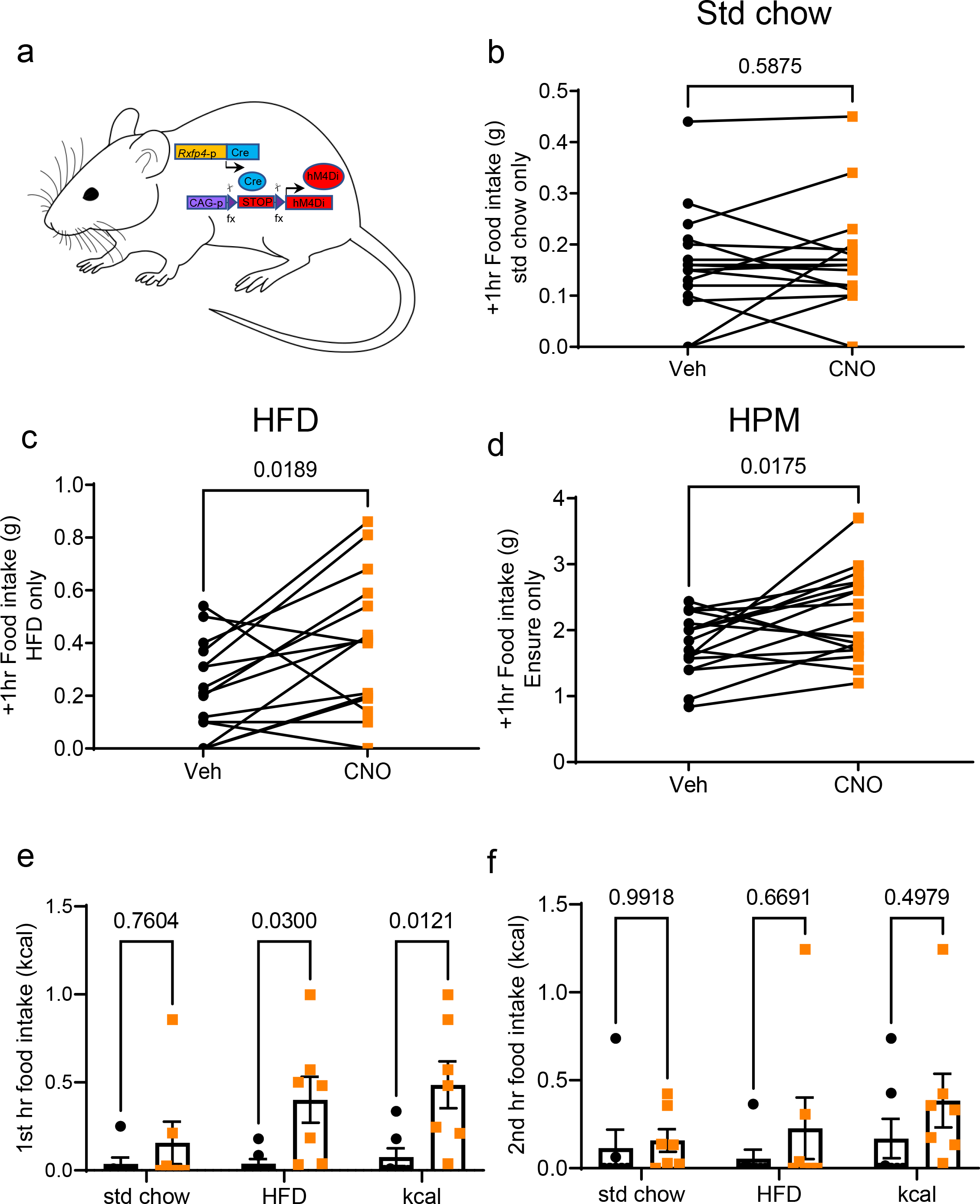
CNO alters feeding and food preference in RXFP4^wb-Di^ mice. a) Schematic of RXFP4^wb-Di^ mouse model. b-d) Food intake in RXFP4^wb-Di^ mice of standard chow (t=0.5536, p = 0.5875) (b) or HFD (45%) (t=2.612, p = 0.019) (c) or liquid Ensure (HPM) (t=2.648, p = 0018) (d) 1 hour post CNO/vehicle treatment at 11:00 (n = 17, paired two-tailed t test, animals adapted to the appearance of test meal over the course of two weeks). (e) 1^st^ hr (effect of treatment [F(1,36)=14.86, p = 0.0005], std chow p = 0.7604, HFD p = 0.03, kcal p = 0.012) and (f) 2^nd^ hr food (effect of treatment [F(1,36)=2.219, p = 0.15], std chow p = 0.9918, HFD p = 0.6691, kcal p = 0.50) consumption after CNO (orange) or saline (black) injection at 11:00 in mice housed in metabolic chambers (n = 7, two-way ANOVA with Sidak’s multiple-comparison test) when mice had a choice between std chow or HFD.

We next investigated the effects of whole-body hM3Dq Cre-reporter activation in *Rxfp4*-expressing cells (RXFP4^wb-Dq^) (Fig. 6a). After a 2 h brief fast, to enhance feeding motivation, activation of Dq in *Rxfp4*-expressing cells at the onset of the dark phase had no measurable effect on the food intake of mice offered only standard chow (Fig. 6b). However, when RXFP4^wb-Dq^ animals were habituated to the appearance of HFD or a HPM for 1 hour at the onset of the dark phase, activation of Dq expressing cells with CNO resulted in a marked reduction in HFD or HPM intake (Fig. 6c,d). These results were consolidated in AL fed animals tested in metabolic cages with parallel access to standard chow and HFD. RXFP4^wb-Dq^ activation in AL fed animals at the onset of the dark phase had no effect on standard chow intake, but significantly and transiently reduced HFD consumption (Fig. 6e,f). RXFP4^wb-Dq^ activation also attenuated the increase in energy expenditure associated with the onset of the dark phase, however, there was no effect on ambulatory activity (Suppl. Fig. 4c,d).

**Figure 6:**
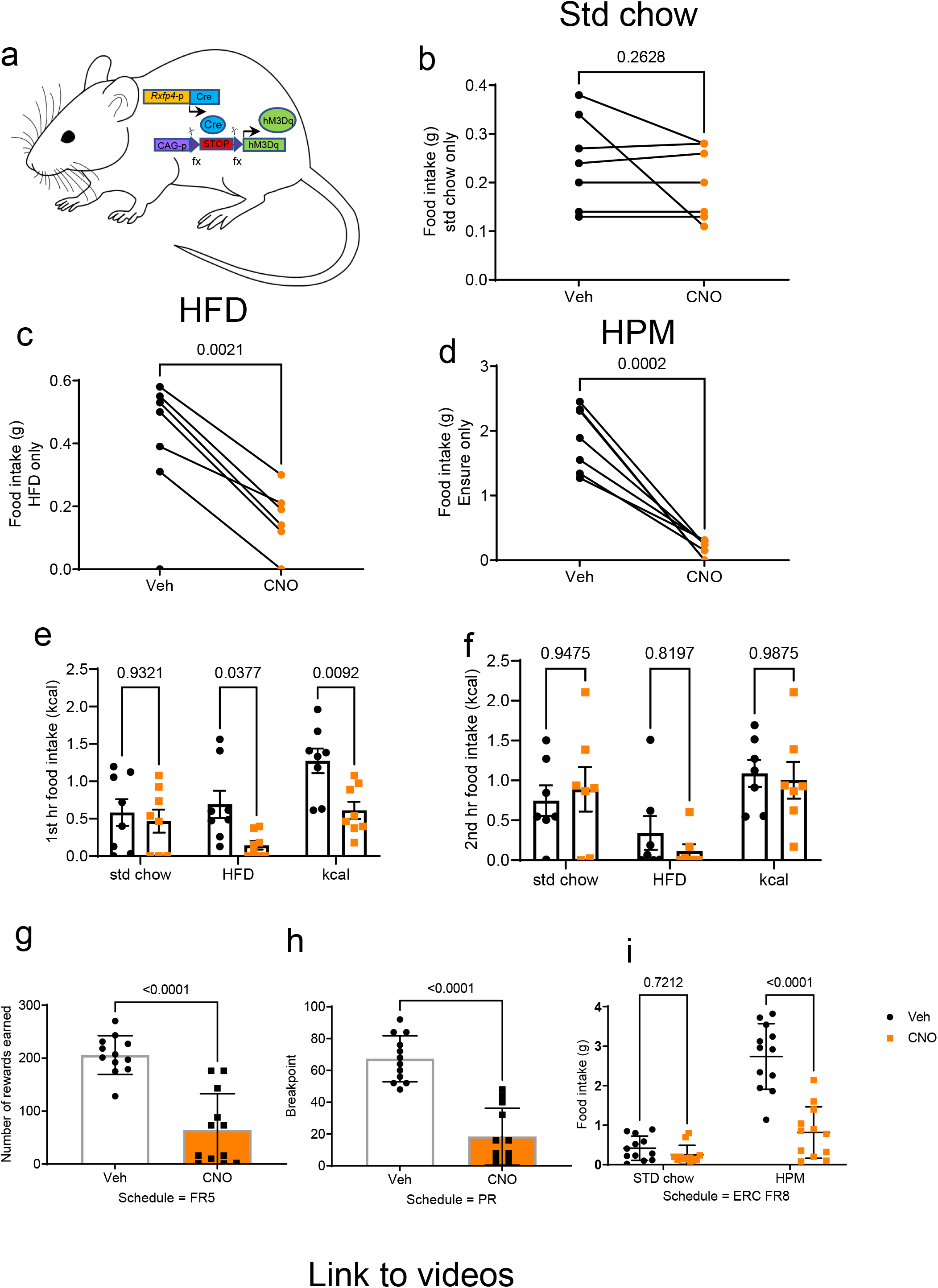
CNO alters feeding and food preference in RXFP4^wb-Dq^ mice. a) Schematic of RXFP4^wb-Dq^ mouse model. b-d) Food intake of (b) standard chow (t=0.1.235, p = 0.2628) (c) HFD (45%) (t=5.136, p = 0.0021) and (d) liquid Ensure (HPM) (t=7.725, p = 0.0002) 1 hour post CNO/saline treatment at 19:00 (n = 7, paired two-tailed t test, animals adapted to the appearance of test meal over the course of two weeks). e) 1^st^ hr (effect of treatment [F(1,42) = 13.17, p = 0.0008, standard chow p = 0.9321, HFD p = 0.0377, kcal p = 0.009) and f) 2^nd^ hr (effect of treatment [F(1,36)=0.1177, p = 0.7335], std chow p = 0.9475, HFD p = 0.8197, kcal p = 0.9875) food consumption after CNO (orange) or saline (black) injection at 19:00 in mice housed in metabolic chambers (n = 7, two-way ANOVA with Sidak’s multiple-comparison test) when mice had access to a choice between std chow and a HFD. g-i) Performance parameters in mice trained in operant chambers when treated with CNO (orange) or vehicle (black): g) Number of rewards earned in FR5 (n = 12, t=6.874, p < 0.001, paired two-tailed t test). h) Breakpoint in PR4 (number of target responses emitted by an animal in the last successfully completed trial, before session termination or 60 min time-out) (n = 12, t=9.357, *** p < 0.001, paired two tailed t test). i) ERC performance post vehicle (black) or CNO (orange) treatment between 11-13:00 (n = 12, effect of treatment [F(1,44)=41.38, p < 0.0001, std chow p = 0.72, HPM p < 0.0001, two-way ANOVA with Sidak’s multiple-comparison test, animals calorically restricted to 95% BW).

To probe whether *Rxfp4*-expressing cells play a role in the motivational aspects of feeding, we calorically restricted male RXFP4^wb-Dq^ animals to 95% body weight and placed them in operant chambers. Mice were tested with a fixed ratio (FR) schedule, requiring 5 nose pokes to release a food reward (liquid Ensure), or a progressive ratio (PR) schedule requiring increasing number of nose pokes for each subsequently earned reward (in this case, +4, i.e. 1, 5, 9, 13, etc). RXFP4^wb-Dq^ mice treated with CNO completed fewer attempts under FR to earn individual Ensure rewards (Fig. 6g). Under a PR schedule, they exhibited a reduced breakpoint, i.e. CNO-treated mice stopped working for the HPM-reward at lower ratios than when receiving vehicle treatment (Fig. 6h). In an effort related choice (ERC) paradigm, where animals had the choice of working for a HPM (FR8, liquid Ensure) or consuming freely available standard chow, CNO treatment reduced HPM consumption (Fig. 6i). However, animals consumed similar amounts of standard chow and displayed otherwise normal behaviour (supplementary video 1), suggesting that activation of *Rxfp4*-expressing cells reduced motivation for the HPM rather than inducing generalised malaise.

To assess whether the *Rxfp4*-expressing population in the VMH is involved in the feeding phenotype observed in RXFP4^wb-Dq^ mice, the effect of acute chemogenetic manipulation of RXFP4^VMH^ neuron activity on food intake was investigated. Male and female *Rxfp4*^GCAMP3^ mice received bilateral VMH injections of Cre-dependent hM3Dq-expressing rAAVs (rAAV-hSyn-DIO-hM3D(G)q-mCherry) designed to preferentially target neurons, to produce RXFP4^VMHDq^ mice (Fig. 7a). Targeting efficiency was subsequently determined by immunohistochemistry (Fig. 7b), or in a subset by fluorescent microscopy in live slices (see below). All mice demonstrated robust transduction that was limited to the target region. To confirm the functional activation of these RXFP4^VMHDq^ neurons, we generated ex vivo brain slices from a subset of RXFP4^VMHDq^ mice (Fig. 7c). Calcium imaging demonstrated that ex vivo treatment with CNO activated mCherry/GCaMP3 positive RXFP4^VMHDq^, stimulating an increase in the frequency of calcium oscillations and an increase in GCaMP3 fluorescence in *Rxfp4*-expressing somas (Fig. 7c,d).

**Figure 7:**
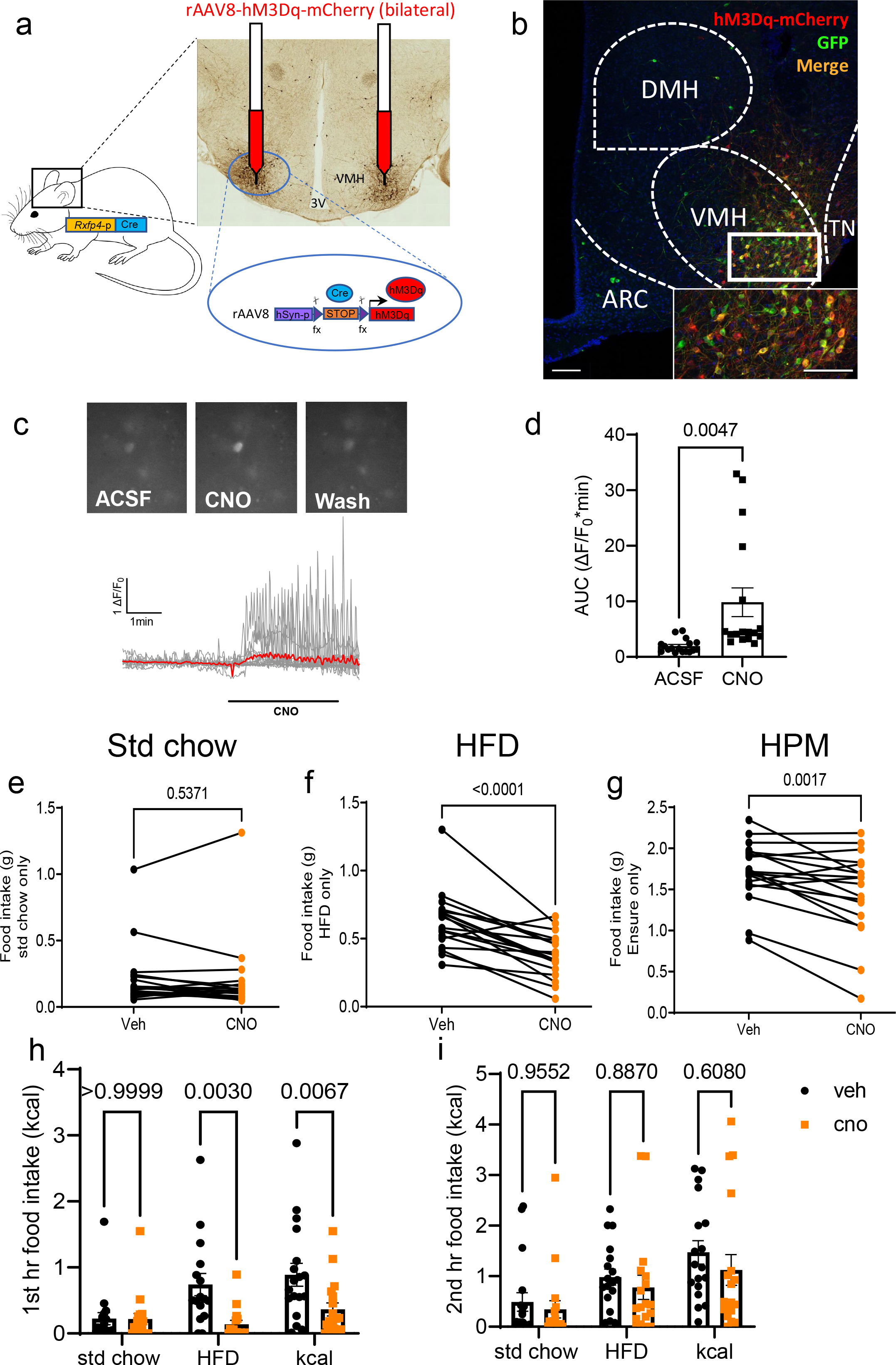
Ventromedial hypothalamic *Rxfp4*-expressing neurons regulate food preference. a) rAAV-hM3Dq-mCherry was bilaterally injected into the ventromedial hypothalamus of *Rxfp4*-Cre mice. Targeting efficiency was confirmed post perfusion fixation. Representative image showing ARC, DMH, VMH and TN with only VMH demonstrating rAAV-hM3Dq-mCherry expression. c) Representative images of RXFP4^VMHDq^ cells before, during and after bath applied CNO (15 µM) showing somatic GCaMP3 fluorescence (top panel), with traces from individual cells (bottom panel, mean intensity shown in red). d) Quantification of responses showing median AUC before and during bath applied CNO (t= 3.285, p = 0.0047 paired two-tailed t test, n = 17 cells from 3 mice). e-f) Food intake of (e) standard chow (t=0.63, p = 0.5371) (f) HFD (45%) (t=5.630, p < 0.0001) and (g) liquid Ensure (HPM) (t=3.714, 0.0017) 1 hour post CNO (orange) or vehicle (black) treatment at 19:00 (n = 18, paired two-tailed t test, animals adapted to the appearance of test meal over the course of two weeks. (h) 1^st^ hr (effect of treatment [F(1,98)=14.70, p = 0.0002, std chow p > 0.9999, HFD p = 0.003, kcal p = 0.0067) and (i) 2^nd^ hr (effect of treatment [F(1,102)=1.65, p = 0.2014, std chow p = 0.9552, HFD p = 0.887, kcal p = 0.6080) food consumption of RXFP4^VMHDq^ mice in metabolic chambers, when mice had access to a choice between std chow and a HFD, post CNO (orange) or vehicle (black) treatment (n = 18 per group, mean ± SEM). Scale bars = 100µm.

The effect of chemogenetic activation of this cell population in vivo on food intake was studied in a crossover design. In chow fed mice, and in line with RXFP4^wb-Dq^ animals, CNO treatment of RXFP4^VMHDq^ mice had no effect on standard chow intake at the onset of the dark phase (Fig. 7e). When animals were habituated to the appearance of a HFD or HPM at the onset of the dark phase, CNO resulted in a significant reduction in food intake (Fig. 7f,g). When offered HFD and chow diet in parallel in metabolic cages, CNO significantly reduced intake of the HFD whilst intake of standard chow was not altered, resulting in an overall reduced caloric intake (Fig. 7h). As seen with RXFP4^wb-Dq^ animals, this was a transient effect no longer apparent in the 2^nd^ hour post CNO administration (Fig. 7i). CNO had no effect on energy expenditure or ambulatory activity in these RXFP4^VMHDq^ animals (Suppl. Fig. 4e,f).

## Discussion

The *Rxfp4*-Cre mouse model generated in this study has enabled the identification of central *Rxpf4*-expressing cells, and aided the transcriptomic, functional and anatomical characterisation of a hypothalamic *Rxfp4*-expressing neuronal population. We show that *Rxfp4* is expressed in key feeding centres of the brain, including neurons of the VMH. Ex vivo CAMPER imaging revealed that INSL5 reduces cAMP levels in *Rxfp4*-expressing VMH neurons and central administration of INSL5 into the VMH increased HFD and HPM intake in lean AL fed mice in vivo. Mimicking the native physiology of INSL5-RXFP4 via Di recapitulated these results, whereas chemogenetic cell activation via Dq suppressed HFD and HPM intake. Selective chemogenetic activation of *Rxfp4*-expressing cells in the VMH alone (targeted via rAAV-Dq) also suppressed HFD and HPM intake, reflecting their position in brain circuits implicated in homeostatic and hedonic regulation of food intake(20). Ablation of the VMH *Rxfp4*-positive cell population resulted in lower HFD consumption and body weight, mirroring the phenotype of global *Rxfp4* knockout mice (4) and supporting the conclusion that *Rxfp4* neurons in the VMH play a role in the physiological drive to consume highly palatable foods. Overall, these data identify hypothalamic RXFP4 signalling as a key regulator of food intake and preference.

*Rxfp4* expression has been difficult to localise due to low mRNA expression levels and the lack of suitable verified antibodies. *Rxfp4* expression was previously reported in the colon (4) and in enteroendocrine tumor cell lines (21, 22). In contrast to our previous report (4), we detected *Rxfp4* mRNA in the hypothalamus (Fig. 1g,h) and found substantial *Rxfp4*-dependent reporter expression in multiple brain regions, with distinct *Rxfp4*-expressing cell populations in the accessory olfactory bulb, RSC, VMHvl, and mammillary body (Suppl. Fig. 1). While it could be argued that this reflects lineage tracing from *Rxfp4*-positive precursor cells, the detection of *Rxfp4* mRNA by RNAscope, the activation of Cre-dependent rAAV-constructs when stereotactically injected into the adult VMH, the cAMP responsiveness of *Rxfp4*-positive cells to locally-perfused INSL5 in slice preparations, and the effect of stereotactically injected INSL5 on feeding behaviour, confirm active *Rxfp4*-promoter activity and consolidate active RXFP4 expression and function in the adult mouse brain (Fig. 1g,h, Fig. 7b).

To characterise the transcriptomic profile of hypothalamic *Rxfp4*-expressing cells we performed scRNA-Seq. Although some cells will have been lost and some genes may have exhibited altered expression during the cell dissociation and sorting process, the results allowed us to cluster *Rxfp4*-expressing cells into several subpopulations each characterised by a profile of cell-specific marker genes (Fig. 2a,b). *Rxfp4* was identified in microglia, ependymocytes and endothelial cells that potentially constitute the blood brain barrier, suggesting that INSL5 may additionally exert effects on non-neuronal cells(23). Clustering of the *Rxfp4*-expressing neuronal population indicated a predominance of glutamatergic neurons with few GABAergic clusters (Fig 2d). Cluster 6 is of note given the expression of *Glp1r*, *Cckbr*, *Sstr1* and *Sstr2*, suggesting an overlap with other known appetite-modulating gut peptide receptors. The cocaine- and amphetamine-regulated transcript (*Cartpt*), expressed in cluster 3, and cannabinoid receptor 1 (*Cnr1*), expressed in clusters 1, 4 and 5, (Fig 2d, Suppl. Fig. 3) have also been implicated in energy homeostasis(24, 25). Cluster 1 displayed markers of a VMHvl *Esr1* population (*Esr1, Pgr, Tac1, Cckar, Rprm* and *Oxtr*) previously associated with food intake and energy expenditure(12, 15, 16). However, chemogenetic activation of RXFP4^VMHDq^ neurons did not result in increased energy expenditure or ambulatory activity, contrasting with the previously described Esr1^VMHDq^ phenotype(16). The role of RXFP4^VMH^ cells in feeding regulation is further suggested by the co-expression of the neuropeptide receptors *Mc4r* and *Nmur2.* MC4R activation has been linked to suppressed food intake through regulation of *Bdnf* expression in the VMH (26) – *Bdnf* was co-expressed in cluster 1 neurons in our dataset (Fig. 2d). Acute administration of NMUR2 agonists have been shown to decrease feeding, with one agonist being somewhat selective to HFD intake regulation (27), mirroring RXFP4^VMHDq^ cell activation (Fig 7).

Mapping of the retrograde inputs to and anterograde projections from the *Rxfp4*-expressing cells in the VMH revealed a distinct neural circuitry surrounding this hypothalamic population. While we aimed to target the VMHvl specifically during stereotactic injections, disparities between current mouse brain atlases make it difficult to distinguish the VMHvl from the adjacent VMHc and TN, which have also been implicated in feeding behaviour (13, 30) . It is therefore possible that some *Rxfp4*-expressing cells in the VMHc and TN were transfected by the viral vectors and acted as starter cells in these tracing experiments. The *Rxfp4*-expressing population we targeted is therefore best described as RXFP4^VMH^. Monosynaptic inputs to RXFP4^VMH^ cells were labelled predominantly from brain regions involved in homeostatic regulation of food intake such as the ARC, PVH and LHA (Fig. 3f). Within the ARC, two populations of neurons have been intensely studied with regards to feeding regulation: pro-opiomelanocortin (POMC)/cocaine- and amphetamine-regulated transcript (CART)-expressing neurons inhibit food intake while Agouti-Related Peptide (AgRP)/neuropeptide Y (NPY)-expressing neurons stimulate food intake. These neurons integrate nutritional and hormonal signals from the periphery and send projections to multiple brain regions including the VMH (31). Although the VMH is not thought to be the main target of arcuate POMC projections (32), the co-expression of *Mc4r* in the RXFP4^VMH^ neurons suggests that POMC neurons may be part of the ARC innervation of RXFP4^VMH^ neurons.

RXFP4^VMH^ cells predominantly project onto regions associated with reward and motivation-related behaviours (Fig. 3c, Suppl. Fig. 3) such as the BNST, POA, CeA, paraventricular thalamic nucleus (PVT) and ventral tegmental area (VTA) (33–35), potentially underlying our finding that chemogenetic activation of *Rxfp4*-expressing cells in RXFP4^wb-Dq^ mice reduced an animal’s drive to seek out and work for a highly palatable food reward (Fig 6g-i). Taken together, these data suggest that *Rxfp4*-expressing cells may influence motivation and reward-related behaviour via regulation of central reward signalling pathways. RXFP4^VMH^ cells also send projections to the PAG and PBN, two integration sites responsible for relaying sensory information between the forebrain and hindbrain and coordinating behaviour in response to various stimuli including metabolic, gustatory and nociceptive inputs (36–38). This RXFP4^VMH^ projection map aligns with previously identified projection regions from the VMHvl and SST-expressing cells in the TN (13, 39). Interestingly, all retrograde-labelled input regions also received projections from RXFP4^VMH^ cells suggesting a high level of bidirectional connectivity within the RXFP4^VMH^ signalling network (Fig. 3g). Similar bidirectional connectivity has been shown for an *Esr1*+ve VMHvl population (39). The RXFP4^VMH^ neural network established in the present study suggests these cells may integrate metabolic and nutritional cues either directly or via other hypothalamic regions and regulate the reward system to influence ingestive behaviours. The low number of RXFP4^VMH^ input regions compared to projection regions suggests these cells may comprise an early node in this network.

Inhibition of *Rxfp4*-expressing cells via direct intra-VMH infusion of INSL5 or whole body Di acutely increased intake of both a HFD and a HPM in the home cage and when animals were offered a choice of standard laboratory chow and HFD in metabolic cages, without altering energy expenditure and activity in both male and female mice (Fig 1j,k, Fig 5, Suppl. Fig 4a,b). This is consistent with our previous demonstration that INSL5 administration increased food intake (4, 5). Activation of Di receptors should, at least in part, mimic the Gαi/o coupling of RXFP4 (2). By contrast, activation of *Rxfp4*-expressing cells with Dq produced a robust suppression of HFD and HPM intake (Fig 6). Targeting the RXFP4^VMH^ population with an rAAV-Dq reporter recapitulated the findings with the whole body-Dq approach (Fig. 7). Whilst any of the identified *Rxfp4*-expression sites could participate in the feeding phenotype of the global DREADD reporter mice, the VMH-specific rAAV-Dq reporter phenotype indicates that this hypothalamic population either underlies or at least contributes to the observed anorexigenic effects in the RXFP4^wb-Dq^ model.

Our data suggest that activation of *Rxfp4*-expressing cells in the VMH suppresses the consumption and drive to work for calorie dense HFD and HPM. The importance of the VMH in the regulation of feeding and metabolism has been disputed (reviewed in (40)). Initial studies suggested the VMH might be a “satiety centre”, as VMH-lesioned rats, particularly females, over-consumed a HFD when fed AL (41), despite being seemingly less willing to work for food on a fixed ratio lever-pressing paradigm (42). The observed hyperphagia has subsequently been linked to additional damage to adjacent hypothalamic structures (see King 2006(40) for discussion), while the reduced motivation was not observed when rats were trained pre-operatively, suggesting that the VMH lesion may have altered “trainability” rather than feeding motivation (40, 43). Lesioning studies, whilst informative, lack cellular precision and damage neural connections to other feeding centres within the brain. More recent work employing immunohistochemistry, RNA sequencing, chemogenetics and neuronal projection mapping, has demonstrated that the VMH consists of anatomical subdivisions made up of distinct cell populations(15, 44, 45). Functional studies suggest neurons in the central and dorsomedial VMH regulate feeding, energy expenditure and glucose homeostasis (30, 46), while the VMHvl is more frequently implicated in the control of social and sexual behaviours(45, 47, 48). However, several studies have demonstrated the involvement of VMHvl neurons in energy expenditure and feeding behaviour (12, 49). Increases in physical activity have been observed following chemogenetic activation of NK2 homeobox transcription factor 1 (*Nkx2-1*)-expressing neurons in the VMHvl of female rats, while knockout of *Nkx2-1* in the VMHvl leads to decreased physical activity and thermogenesis (50). Furthermore, chemogenetic activation of *Esr1*-expressing VMHvl neurons was found to stimulate physical activity and thermogenesis in both sexes (16). *Esr1* signalling in the VMHvl was previously demonstrated to influence food intake, energy expenditure and glucose tolerance as VMHvl-restricted knockdown of *Esr1* resulted in increased food intake, decreased physical activity and thermogenesis, reduced glucose tolerance and obesity in female rats (12). Activation of RXFP4^VMHDq^ neurons reduced HFD and HPM intake but had no effect on chow intake, energy expenditure or ambulatory activity (Fig. 7, Suppl. Fig. 4e,f). This, alongside the transcriptomic profile and neural circuitry of RXFP4^VMH^ neurons, suggests that these neurons comprise a distinct VMH population modulating food intake and preference based on the rewarding aspects of food rather than the homeostatic food intake or energy expenditure responses observed during chemogenetic manipulation of other VMH populations.

We recognise several limitations to this study. First, the exact classification of *Rxfp4*-expressing brain regions is difficult given the disparities between current brain atlases. Whilst we aimed to target *Rxfp4*-expressing cells in the VMHvl, the inexact nature of stereotactic injections may have resulted in additional targeting of neurons in adjacent VMHc, LHA and TN regions, which may have contributed to the food intake phenotype of RXFP4^VMHDq^ mice. The fact that only a subset of Rxfp4^CAMPER^ neurons responded to INSL5 in acute slices likely reflects technical limitations of this preparation. The similarities of the feeding outcomes of intraparenchymal INSL5 in WT mice and CNO application in Rxfp4^VMHDq^ mice together with RNAscope confirmation of Rxfp4-Cre labelled VMH neurons makes the alternative explanation of substantial additional off target Cre-expression unlikely, but we cannot fully rule this out. It is also uncertain whether INSL5 is the endogenous ligand acting on *Rxfp4*-expressing cells in the brain. We have previously been unable to identify *Insl5*-expressing cells in the mouse brain (5) and there is no evidence that INSL5 can cross the blood brain barrier. In addition, relaxin-3 also activates RXFP4 (2), is expressed in the mouse brain and is orexigenic (51), hence it is possible that relaxin-3 is involved in central RXFP4 action. The viral tracing techniques used to decipher the RXFP4^VMH^ neuronal network also have some limitations. In the retrograde tracing experiments, it was difficult to detect rAAV-GFP immunoreactive cells, making it hard to confirm the exact starter cells in this experiment. Furthermore, while the ChR2-mCherry construct is preferentially targeted to axon terminals, it is possible that some of the mCherry-labelled fibres in adjacent regions, such as the ARC, are in fact dendrites (52) which may underlie our inability to detect retrograde-only labelled regions. Nevertheless, we have been able to identify distinct regions that project onto and receive projections from *Rxfp4*-expressing cells in the VMH that connect these cells to known feeding-related neural networks. Finally, it is difficult to explain apparently similar outcomes of acute chemogenetic activation and chronic ablation of RXFP4^VMH^ neurons, both resulting in reduced HFD consumption. The chronic ablation outcome would be compatible with a positive valence of Rxfp4^VMH^ neurons, whilst the acute Dq experiment would be compatible with a negative valence of the same neurons on HFD consumption, and both interpretations would be compatible with the response to acute parenchymal Insl5 and the phenotype of Rxfp4-knock-out mice. Whilst we are currently unable to distinguish these two interpretations, we nevertheless establish Rxfp4^VMH^ neurons as integral players in a complex neural network consuming highly palatable food consumption.

In summary, we have characterised a previously unrecognised population of ventromedial hypothalamic cells that express *Rxfp4* in mice, identified projections in homeostatic and hedonic feeding centres in the CNS, and demonstrated that acute manipulation of these cells modulates HFD/HPM intake without affecting chow intake or energy expenditure. Together, these findings suggest *Rxfp4*-expressing neurons in the VMH are key regulators of food preference and represent a target for the modulation of feeding behaviour.

## Methods

### Animals

All experiments were performed under the UK Home Office project licences 70/7824 and PE50F6065 in accordance with the UK Animals (Scientific Procedures) Act, 1986 and approved by the University of Cambridge Animal Welfare and Ethical Review Body. All mice were from a C57BL/6 background and were group-housed and maintained in individual ventilated cages with standard bedding and enrichment in a temperature and humidity controlled room on a 12h light:dark cycle (lights on 7:00) with *ad libitum* (AL) access to food (Scientific Animal Food Engineering) and water unless otherwise stated. Groups were randomised by body weight and the researcher was blinded to treatment.

### Mouse models

To express Cre recombinase under the control of the *Rxfp4* promoter, we replaced the sequence between the start codon and the stop codon in the single coding exon of *Rxfp4* in the murine-based BAC RP24-72I4 (Children’s Hospital Oakland Research Institute) with iCre (53) sequence using Red/ET recombination technology (GeneBridges) (Fig 1A). The resulting BAC was used to create BAC-transgenic mice – of four initial founders, two passed the transgene to their offspring; both resulting lines showed similar Cre-reporter expression and one line, Rxfp4-73, was used throughout this manuscript (see Suppl. methods 1 for further information). Several Cre-reporter transgenes, in which expression is only activated after removal of a fxSTOPfx cassette, were used, resulting in expression of EYFP (54), Dq, Di (55), GCaMP3 (56), or tdRFP (57), respectively.

### Viral injections

Viral injections were performed in male and female *Rxfp4*-Cre mice aged between 9 and 16 weeks. The surgical procedure was performed under isoflurane anaesthesia, with all animals receiving Metacam prior to surgery. Mice were stereotactically implanted with a guide cannula (Plastics One) positioned 1mm above the VMH (A/P: -1.7 mm, D/V: -4.5 mm, M/L: +/-0.75 mm from bregma). Bevelled stainless steel injectors (33 gauge, Plastics One) extending 1mm from the tip of the guide were used for injections. For anterograde viral tracing experiments, 200 nL rAAV-DIO-ChR2-mCherry (Addgene, 20297-rAAV8, 1.9 × 10^1^³ vg/ml) was injected unilaterally at 75 nL/min. Mice were culled three weeks after injection. For retrograde viral tracing experiments, AVV2-FLEX-TVAeGFP-2A-oG (rAAV2-TVAeGFP-oG) and Rabies-ΔG-EnvA-mCherry (Rab-ΔG-EnvA-mCherry) viruses were generated by Ernesto Ciabatti (MRC Laboratory of Molecular Biology, Cambridge). Mice were injected unilaterally with 200 nL AVV2-TVAeGFP-oG (1×10^12^ vg/mL) at 75 nL/min followed by 500 nL Rab-ΔG-EnvA-mCherry (2×10^9^ iu/mL) at 75 nL/min three weeks later. Mice were culled seven days after the second injection. For the RXFP4^VMHKO^ studies, 200 nL ssAAV9-hEF1α-dlox-DTA(rev)-dlox-WPRE-hGHp(A) (rAAV-DTA: ETH VVF, 6.2×10^12^ vg/mL) or AAV2-hSyn-DIO-mCherry (rAAV-mCherry: Addgene, 50459-AAV2, 4×10^12^ vg/mL) was injected bilaterally at 75 nL/min and mice were allowed 2 weeks recovery prior to testing. For phenotyping experiments, 200nL rAAV8-hM3D(G)q-mCherry (Addgene 44361-rAAV8, 4×10^12^ vg/mL) was injected bilaterally at 50nl/min and mice were allowed 2 weeks recovery prior to testing.

### Intra-VMH infusion

Mice were stereotactically implanted with a guide cannula (Plastics One) positioned 1mm above the VMH (A/P: -1.7 mm, D/V: -4.5 mm, M/L: +/-0.75 mm from bregma). Bevelled stainless steel injectors (33 gauge, Plastics One) extending 1mm from the tip of the guide were used for infusions in free-moving mice. An intra-VMH cannula was placed under isoflurane anaesthesia and the animals was given 1 week to recover. Mice were given AL access to standard (std) chow.

### Food intake

Food intake studies were performed in a cross-over design, on age-matched groups, a minimum of 72 hours apart. Mice were administered vehicle or INSL5 (5ug/animal, Phoenix Biotech, 035-40A) at 12:00-15:00 following a 2h fast. Food was weighed 1h post-injection. For experiments assessing the effect of global RXFP4 Di and Dq activation, animals were singly housed prior to the experiment. Mice were administered 1mg/kg CNO (Sigma, C0832) or an equivalent volume of vehicle containing a matched concentration of DMSO (Sigma, D2650). For light phase activation, animals were injected with vehicle or CNO at 11:00 (± 30mins) following a 2h fast. Food was weighed at 1h post-injection. For dark phase activation, animals were injected with vehicle or CNO at 19:00 at the onset of the dark phase following a 2h fast. Food was weighed at 1h post-injection. In trials with a high fat diet (HFD) and a highly palatable meal (HPM, liquid Ensure, Abbott Laboratories, 353-3601), mice were habituated to the appearance of the test meal for 1 hour (5 days per week) for two weeks prior to testing.

### Operant chambers

Twelve male RXFP4^wb-Dq^ mice (weighed 3 times weekly) were food restricted to maintain 95% body weight for two weeks prior to training and testing in standard mouse Bussey-Saksida touchscreen chambers (Campden Instruments Ltd, Loughborough, UK). Training and testing procedures were conducted as previously described (58). Briefly, mice were trained to touch a screen for a reward (the HPM, 20 µL) under a fixed ratio (FR) schedule for 2 weeks, progressing from FR1 to FR5 (training deemed successful when the animal earnt 30 rewards within 1 hour), followed by testing. Mice then progressed to progressive ratio (PR, increment +4 i.e. 1, 5, 9, 13, etc), where the breakpoint was defined as the last reward earned before 5 minutes elapsed without operant response. Following testing of the breakpoint, mice progressed to the effort related choice schedule (ERC) – mice were trained on FR8, with the addition of standard chow to the operant arena. Once animals successfully earned 30 rewards within 60 minutes, testing was undertaken. The 60-minute training and testing sessions took place at the same time each day (between 10:00-13:00).

### Metabolic cages

Animals were acclimated to metabolic cages prior to study and data collection. Oxygen consumption and carbon dioxide production were determined using an indirect calorimetry system (Promethion, Sable Systems, Las Vegas, NV). The system consisted of 8 metabolic cages (similar to home cages), equipped with water bottles and food hoppers connected to load cells for continuous monitoring, in a temperature and humidity-controlled cabinet. The respiratory exchange ratio (RER) and energy expenditure (via the Weir equation) were calculated, whilst ambulatory activity was determined simultaneously. Raw data were processed using ExpeData (Sable Systems). Animals were exposed to standard chow and a HFD during metabolic assessment or standard chow.

### CAMPER and calcium imaging

Mice were anesthetized using sodium pentobarbital at a dose of 180mg/kg (Dolethal, Vetoquinone). A laparotomy was then performed, the heart exposed and the animals were intracardiacally perfused with oxygenated High-Mg^2+^/low Ca^2+^ ice-cold artificial cerebrospinal fluid (perfusion solution; composition in mM: 2.5 KCl, 200 sucrose, 28 _NaHCO3_, 1.25 NaH_2_PO4, 8 Glucose, 7MgCl_2_, 0.5 CaCl_2_; pH 7.4). The brain was extracted from the skull and immersed in oxygenated perfusion solution. 250 μm thick slices of the hypothalamus containing the VMH were cut on a Vibratome (Leica) and placed in a recovery solution (in mM: 3 KCl, 118 NaCl, 25 NaHCO_3_, 1.2 NaH_2_PO_4_, 2.5 Glucose, 7 MgCl_2_, 0.5 CaCl_2_; pH 7.4) at 34 °C for 30 min. Sections were then transferred to standard ACSF (in mM: 3 KCl, 118 NaCl, 25 NaHCO_3_, 10 Glucose, 1MgCl_2_, CaCl_2_; pH7.4) at room temperature for 1h before recording. All solutions were continuously bubbled with 95% O_2_/5% CO_2_.

Imaging was performed using a Zeiss Axioskop2 FS Plus microscope. Slices were immobilised in a glass bottom chamber using a slice anchor (Warner instruments) and imaged using a 40x water immersion objective. For CAMPER imaging, RXFP4^CAMPER^ neurons of the VMH were excited using a LED light source (Dual Optoled, Cairn Research) at 435 for 50ms every 3 seconds and CFP and YFP emission were simultaneously recorded after passing through an optosplit (Cairn; 465-505 and 525-600 nm, respectively). . . For Ca^2+^ imaging, RXFP4^GCaMP3^ neurons were excited at 470 nm for 75 ms every 2 s and emission from 500-550 nm was captured. Images were captured using a charged-coupled camera (Evolve 512 Delta, Photometrics), acquisitions were saved and data logged using Metamorph software (Molecular Devices). Slices were continuously superfused with room temperature standard ACSF at a flow rate of 1 mL per minute. All drugs were added directly in standard ACSF. CNO (15 μM, C0832), IBMX (100 uM, I7018) and forskolin (10 uM, F6886) were purchased from Sigma. INSL5 (100 nM, 035-40A) was purchased from Phoenix Biotech.

Regions of interest (ROIs) and an area determined for background fluorescence were outlined for each image and the mean pixel intensity extracted for each ROI. The mean intensity of the background was subtracted from each ROI. For CAMPER imaging, the fluorescence energy transfer ratio (ΔFRET) of the mTurquoise FRET donor (CFP) and Venus FRET acceptor (YFP) was calculated and is presented on a reverse axis. INSL5 responses were measured by calculating the percentage change in ΔFRET between the maximum IMBX response and the maximum INSL5 response. For Ca^2+^ imaging, fluorescence intensity of the GCaMP3 is presented as ΔF/F0 with F0 being the mean intensity for 5 min before stimulus and ΔF the fluorescence intensity F minus F0. Responses were observed on raw and ΔF/F0 traces and quantified by calculating areas under the curve (AUCs) over 5 min before the stimulus for the ACSF condition and over 5 minutes after the stimulus for the treated conditions. Cells were considered activated by stimuli if the AUC increased by 25% after treatment compared to the AUC before treatment.

### Immunohistochemistry

Colonic tissues were fixed in 4% paraformaldehyde (PFA), dehydrated in 15% and 30% sucrose, and frozen in OCT embedding media (CellPath, Newtown, U.K.). Cryostat-cut sections (8-10μm) were mounted directly onto poly-L-lysine-covered glass slides (VWR, Leuven, Belgium) by the Institute of Metabolic Science Histology Core. Slides were incubated for 1hr in blocking solution containing 10% goat or donkey serum. Slides were stained overnight at 4°C with primary antisera (table 1) in PBS/0.05% Triton X-100/10 % serum. Slides were washed with blocking solution and incubated with appropriate secondary antisera (donkey or goat Alexa Fluor® 488, 546, 555, 633 or 647; Invitrogen) diluted 1:400 and Hoechst diluted 1:1500 for 1 hr. Control sections were stained with secondary antisera alone. Sections were mounted with Prolong Gold (Life Technologies) or hydromount (National Diagnostics, Atlanta, Georgia, USA) prior to confocal microscopy (Leica TCS SP8 X, Wetzlar, Germany). Quantification of cell number was performed using Leica Application Suite X and Image J.

Brain tissue was collected from perfusion fixed mice as previously described (59). Animals were anaesthetised with dolethal sodium pentobarbital solution at 125 mg/kg ip (in saline) and transcardially perfused with heparinised 0.1M phosphate buffered saline (1xPBS) followed by 4% PFA in PBS. Brains were extracted and post-fixed in 4% PFA for 24 hrs at 4°C then transferred to 30% sucrose solution at 4°C for 48 hrs. Brains were sectioned coronally from olfactory bulb to the spinomedullary junction at 25 μm using a freezing microtome and stored in cryoprotectant medium. For diaminobenzidine (DAB) staining, antigen retrieval was used for all experiments prior to antibody incubation. Sections were incubated in 10 mM sodium citrate (Alfa Aesar, A12274) at 80°C for 10 minutes then washed in PBS. Sections were incubated in 0.5% hydrogen peroxide (Sigma, H1009) in milliQ water for 15 minutes then washed in PBS. Sections were blocked with 5% donkey serum in 0.3% Tween20 (VWR, 437082Q) in PBS (PBST) for 1 hr at room temperature, then incubated with GFP antiserum (1:4000; ab5450, Abcam) in blocking solution overnight at 4°C. After washing in 0.1% PBST, sections were incubated with biotinylated anti-goat IgG (1:400; AP180B, Millipore) in 0.3% PBST for 1.5 hrs at room temperature, followed by a 1 hr incubation with streptavidin conjugated to horseradish peroxidase (Vectastain Elite ABC kit, Vector Laboratories, PK-6100) and developed by DAB substrate (Abcam, ab94665). Sections were washed in PBS, prior to dehydration in ethanol and xylene, then mounting/coverslipping with Pertex mounting medium (Pioneer Research Chemicals Ltd., PRC/R/750). For immunofluorescent staining, slices were washed in PBS, prior to blocking for 1 hr in 5% donkey serum then incubation with primary antisera (table 1) in blocking solution overnight at 4°C. Slices were washed in and incubated with the appropriate secondary antisera (Alexa Fluor® 488 or 555; Invitrogen) diluted 1:500 for 2 hrs at room temperature. Following washing, mounted sections were coverslipped on superfrost slides using Vectashield (Vector Laboratories, H-1400-10). Slides were imaged using an Axio Scan.Z1 slide scanner (Zeiss) and confocal microscope (Leica TCS SP8 X, Wetzlar, Germany) with a 20x or 40x objective as indicated. Images were analysed in Halo Image Analysis Software (Indica Labs) and ImageJ.

**Table 1:**
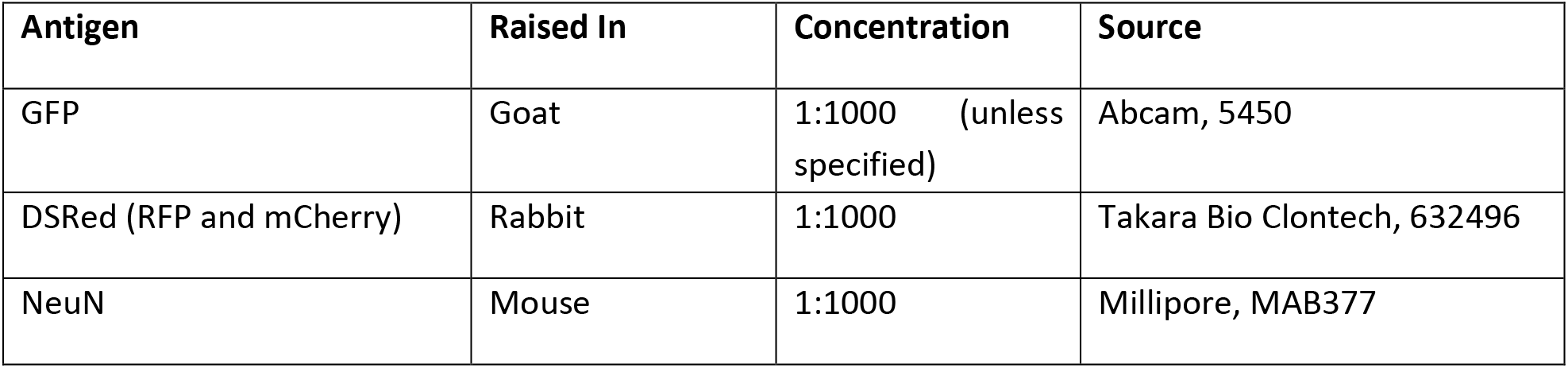
Primary antisera used for immunohistochemistry.

Dissociation, fluorescence-activated cell sorting (FACS) and single cell RNA sequencing

For hypothalamic samples, single cell suspensions were prepared and pooled from twelve female *Rxfp4*-Cre x EYFP mice (biological replicates) as previously described (59). Briefly, mice were sacrificed by cervical dislocation and the hypothalamus dissected into Hibernate-A medium (ThermoFisher, A1247501) supplemented with 0.25% GlutaMAX (ThermoFisher, 35050061) and 2% B27 (ThermoFisher, A1895601). Tissue was digested in Hibernate-A medium without calcium (BrainBits) containing 20 U/mL Papain (Worthington) and 1% GlutaMAX for 30 min at 37°C under agitation (Thermomixer, 500 rpm). After digestion, tissue was triturated in Hibernate-A medium with 3.5 U/mL DNase I (Sigma, D4263). The trituration supernatant was loaded on top of a BSA gradient prepared in Hibernate-A medium, spun for 5 min at 300 rcf, and the pellet was resuspended in Hibernate-A medium + 1% BSA. The cell suspension was filtered through a 70 µm cell strainer into a fresh tube. Fluorescence-activated cell sorting was performed using an Influx Cell Sorter (BD Biosciences, San Jose, CA, USA). Cell gating was set according to cell size (FSC), cell granularity (SSC), FSC pulse-width for singlets, fluorescence at 488 nm/532 nm for EYFP and 647/670 nm for nuclear stain with DraQ5 (Biostatus, Shepshed, Leicester, UK) to exclude cellular debris, aggregates and dead cells. Cells were sorted directly into individual wells of 96-well plates containing lysis buffer. 384 YFP-positive cells were isolated and processed using a Smart-Seq2 protocol (60). Libraries were prepared from ∼150 pg of DNA using the Nextera XT DNA preparation kit (Illumina, FC-131-1096) and Nextera XT 96-Index kit (Illumina, FC-131-1002). Pooled libraries were run on the Illumina HiSeq 4000 at the Cancer Research UK Cambridge Institute Genomics Core. Sequencing reads were trimmed of adapters, aligned to the *Mus musculus* genome (GRCm38, Ensembl annotation version 101) using STAR (v2.7.3a), and raw counts generated using FeatureCounts (Subread v2.0.0). Downstream analyses were performed using the Seurat R package (v4.0.1). Samples were included in the final analyses only if they met all of the following criteria: (a) unique reads > 50%, (b) reads assigned to exons > 20%; (c) number of genes with (FPKM>1) > 3000; (d) total number of unique genes > 200. Genes were included in the final analyses if they were detected in at least 3 samples. Marker genes were identified for clusters using the FindAllMarkers function in the Seurat package; the top 20 gene markers were cross referenced against other bulk and single cell RNAseq databases to assign cell type. Neuronal clusters were identified in a similar fashion using appropriate marker genes.

### RNAscope

Detection of Mouse *Rxfp4* was performed on fixed, frozen sections using Advanced Cell Diagnostics (ACD) RNAscope® 2.5 LS Reagent Kit-RED (Cat No. 322150), RNAscope® LS 2.5 Probe-Mm-Rxfp4 (Cat No. 317588) (ACD, Hayward, CA, USA). To prepare the sections, animals were anaesthetized with sodium pentobarbital solution (50 mg/kg in saline) and transcardially perfused with PBS followed by 4% PFA in PBS. Brains were extracted and post-fixed in 4% PFA for 24 hrs before being transferred to 25% sucrose for 24 hrs at 4°C. Brains were embedded in OCT compound, frozen in a Novec 7000 (Sigma)/dry ice slurry and stored at -80 C. 16μm cryosections containing the hypothalamus were prepared on a Leica CM1950 cryostat (Wetzlar, Germany) at -12°C and stored at -20°C until required.

Slides were thawed at room temperature for 10 min before baking at 60°C for 45 min. The sections were then post-fixed in pre-chilled 4% PFA for 15 min at 4°C, washed in 3 changes of PBS for 5 min each before dehydration through 50%, 70%, 100% and 100% ethanol for 5 min each. The slides were air-dried for 5 min before loading onto a Bond Rx instrument (Leica Biosystems). Slides were prepared using the frozen slide delay prior to pre-treatments using Epitope Retrieval Solution 2 (Cat No. AR9640, Leica Biosystems) at 95°C for 5 min, and ACD Enzyme from the LS Reagent kit at 40°C for 10 minutes. Probe hybridisation and signal amplification were performed according to manufacturer’s instructions, with the exception of increased Amp5 incubation time at 30 min. Fast red detection of mouse *Rxfp4* was performed on the Bond Rx using the Bond Polymer Refine Red Detection Kit (Leica Biosystems, Cat No. DS9390) with an incubation time of 20 min. Slides were then removed from the Bond Rx and were heated at 60°C for 1 hr, dipped in xylene and mounted using EcoMount Mounting Medium (Biocare Medical, CA, USA. Cat No. EM897L). Sections were imaged on a Slide Scanner Axio Scan.Z1 microscope (Zeiss) using a 40x air objective. Three z-stack slices spanning 1.5μM were combined into an extended depth of field image (ZEN 2.6, Zeiss). The CZI files were read into Halo Image Analysis Software (Indica Labs).

### Statistical analysis

Data were plotted using GraphPad Prism 7/8/9 software (GraphPad Software, Inc). Statistical analysis was performed by paired Student’s t-tests, one-way ANOVA with multiple comparisons or two-way ANOVA with multiple comparisons, as indicated. N represents biological replicates. Sample size was computed based on pilot data and previously published data. Data are presented as mean ± SEM and probabilities of p<0.05 were considered statistically significant in all tests.

## Acknowledgements

Metabolic Research Laboratories support was provided by the following core facilities: Disease Model Core, Genomics and Transcriptomics Core, Histology Core, Imaging Core, and Core Biochemical Assay Laboratory (supported by the MRC [MRC_MC_UU_12012/5] and Wellcome Trust [100574/Z/12/Z]). RNA-sequencing was undertaken at the CRUK Cambridge Institute Genomics Core. Cell sorting was performed at the NIHR Cambridge BRC Cell Phenotyping Hub. We thank the Histopathology/ISH Core Facility at Cancer Research UK-Cambridge Institute, for assistance with *in situ* hybridisation. We also would like to thank Chris Riches and Maša Josipović for initial help with metabolic and operant cages. JAT is supported by a NIHR Clinical Lectureship (CL-2019-14-504). Research in the laboratory of F.M.G. and F.R. is supported by the MRC (MRC_MC_UU_12012/3) and Wellcome Trust (220271/Z/20/Z). O.R.M.W. is supported by a BBSRC iCASE PhD studentship partnered with AstraZeneca. F.M.G. and F.R. act as guarantors for this manuscript.

## Author contributions

JEL, ORMW, FMG and FR designed the research studies. JEL, ORMW, DN, CB, LB and AEA conducted experiments. SJK and BG conducted the SmartSeq2 protocol and library preparation for single-cell RNA-sequencing and CAS led the bioinformatic analysis. EC and MT provided the rabies and rAAV viruses for retrograde viral tracing. JAT prepared the tissues and gave guidance for RNAscope. DH and DB co-supervised ORMW and BUP helped with the initial behavioural cage experiments. FR developed the transgenic models. JEL, ORMW, DN, FMG and FR wrote the manuscript. All authors revised and approved the final draft.

## Competing interests

F.M.G. is a paid consultant for Kallyope, New York. The Gribble-Reimann lab currently hosts projects that receive funding from AstraZeneca (O.R.M.W., BBSRC-iCASE), Eli Lilly & Company and LGC. DH and DB are AstraZeneca employees and CB and BUP have also joined AstraZeneca since contributing to this work.

---

**Suppl. Fig 1:**
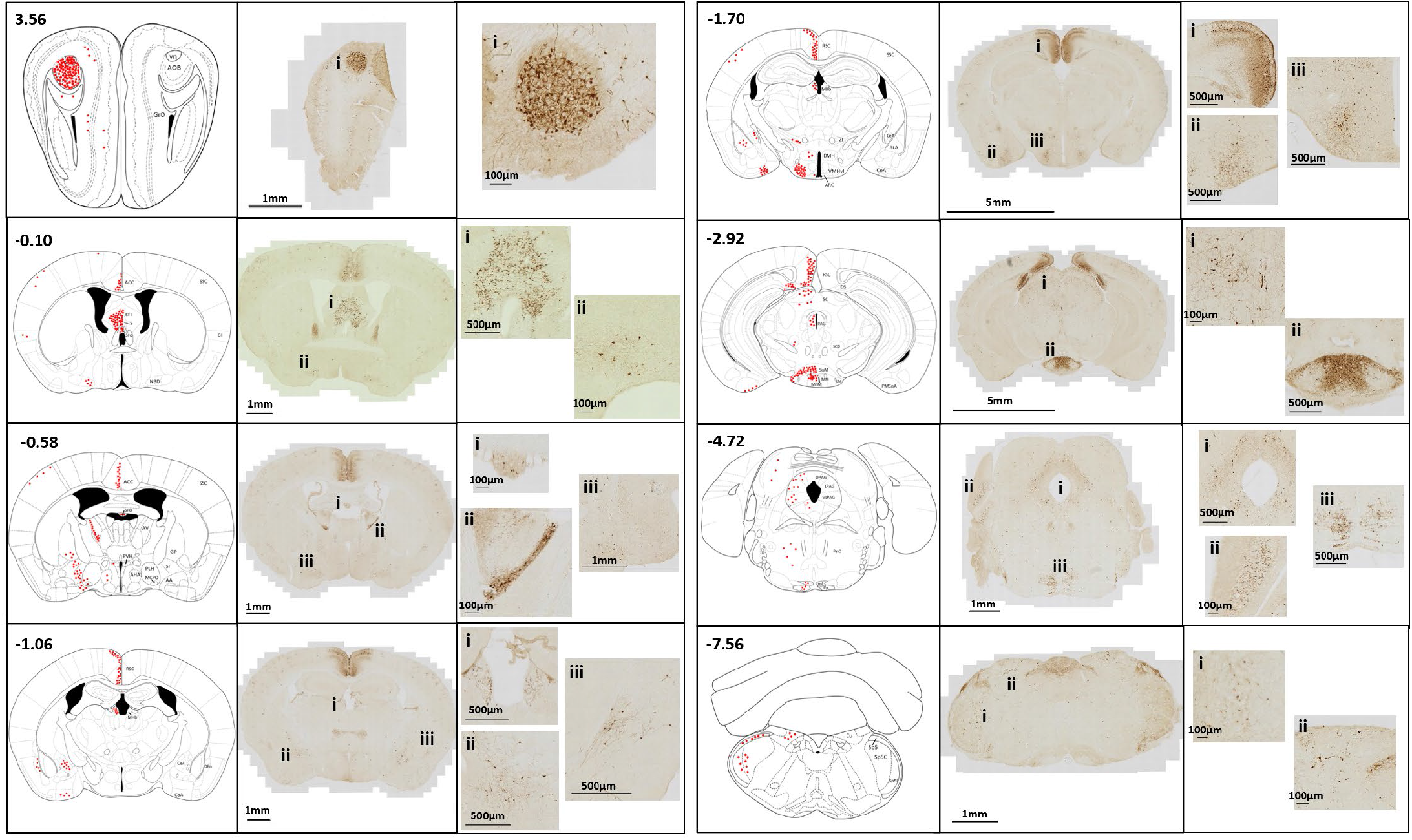
*Rxfp4* expression in the central nervous system. Coronal sections from RXFP4^GCaMP3^ mice stained for GFP immunoreactivity reveal *Rxfp4* expression in various central nuclei. Red circles represent the presence of immunoreactive cells. Reference images based on the Paxinos Mouse Brain Atlas with the A/P coordinates from bregma indicated in the top left corner. AOB: Accesory olfactory bulb; GrO: Granular cell layer of the olfactory; vn: vomeronasal nerve; ACC: anterior cingulate cortex; SSC: somatosensory cortex; GI: granular insular cortex; SFi: septofimbrial nucleus; TS: triangular septal nucleus; SFO: subfornical organ; NBD: nucleus of the diagonal band; AV: anteroventral thalamic nucleus; AHA: anterior hypothalamic area; PLH: peduncular lateral hypothalamus; PVH: paraventricular hypothalamus; MCPO: magnocellular preoptic nucleus; AA: anterior amygdaloid area; SI: substantia innominata; GP: globus pallidus; RSC: retrosplenial cortex; MHb: medial habenular nucleus; CeA: central amygdala; CoA: cortical amygdala; DEn: dorsal endopiriform nucleus; DMH: dorsomedial hypothalamus; VMHvl: ventromedial hypothalamus ventrolateral part; ARC: arcuate nucleus; ZI: zona incerta; BLA: basolateral amygdala; DS: dorsal subiculum; SC: superior colliculus; PAG: periaqueductal grey (D: dorsal, L: lateral, VL: ventrolateral); scp: superior cerebellar peduncle; SuM: supramammillary nucleus; MM: medial mammillary nucleus; MnM: medial mammillary nucleus median part; LM: lateral mammillary nucleus; PMCoA: posteromedial cortical amydala; PnO: pontine reticular nucleus oral part; ml: medial lemniscus; lfp: longitudinal fasciculus of the pons; Cu: cuneate nucleus; Sp5: spinal trigeminal tract (C: caudal part, I: interpolar part).

**Suppl. Fig 2:**
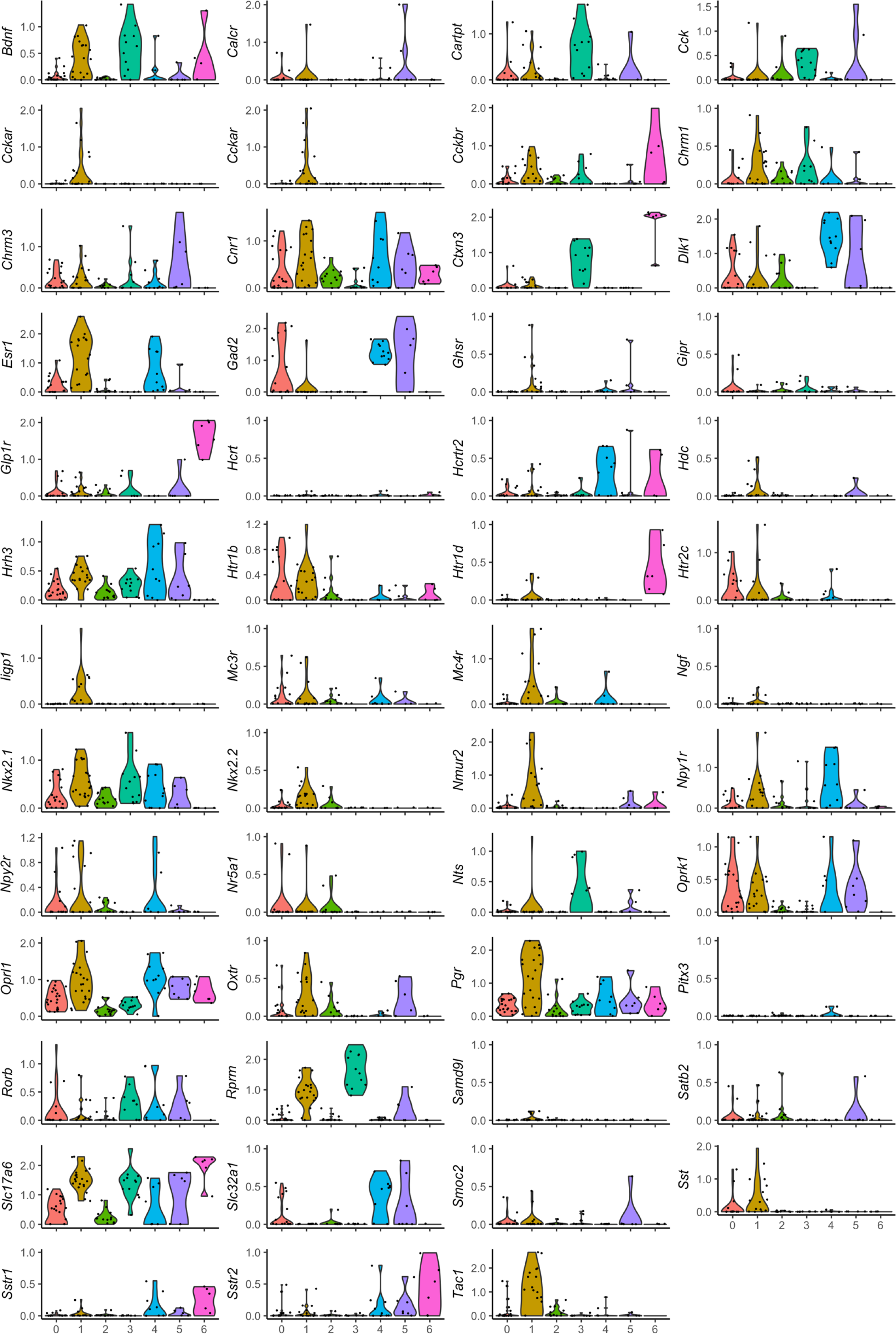
Transcriptomic profiling of the neuronal cluster from scRNAseq of hypothalamic *Rxfp4*-expressing cells. Violin plots showing expression of multiple genes in the hypothalamic *Rxfp4*-expressing neuronal sub-clusters. All gene expression counts are log-normalised with scale-factor = 10^4^.

**Suppl. Fig 3:**
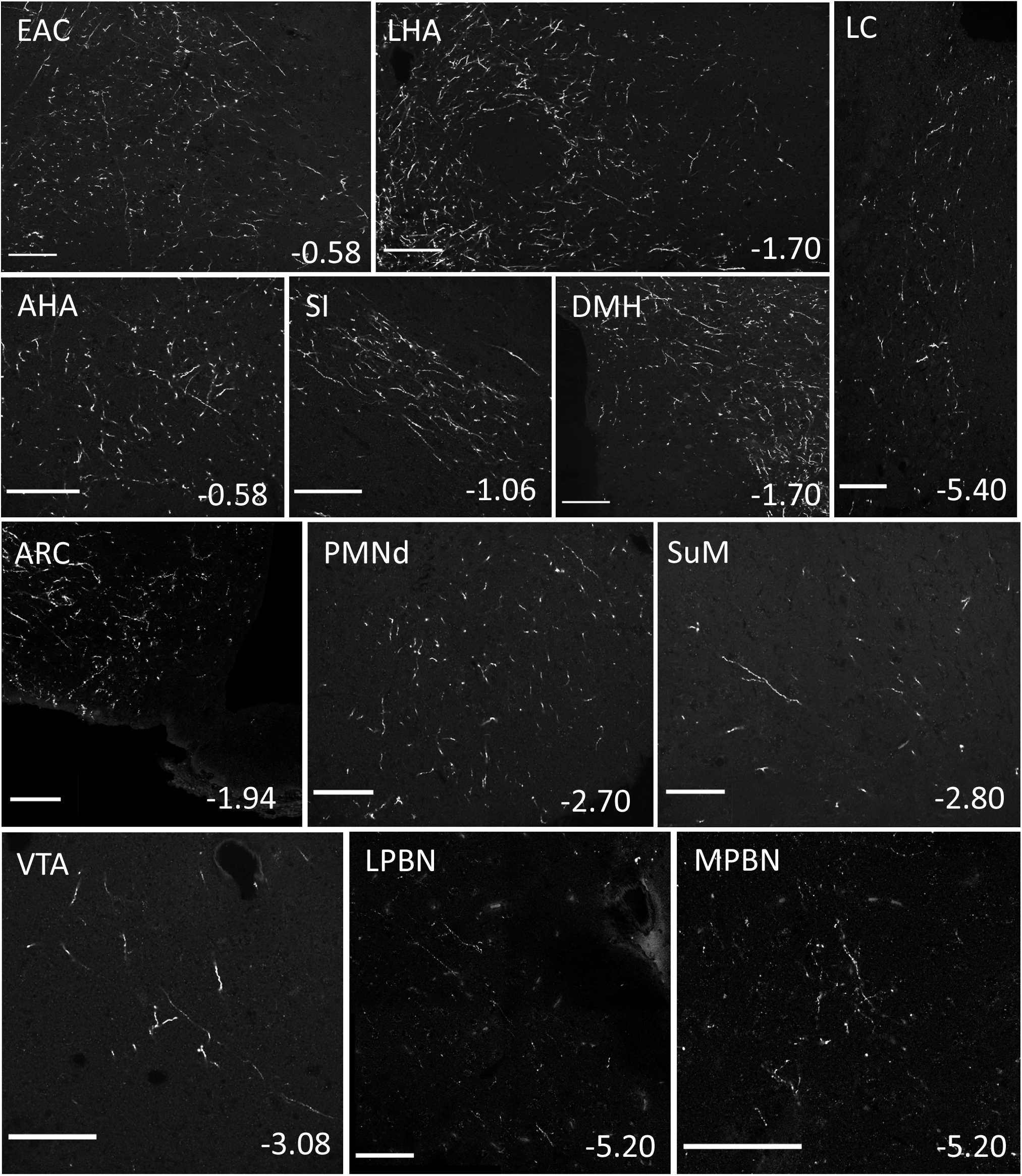
Anterograde projection regions from RXFP4^VMH^ cells. Representative images showing ChR2-mCherry-immunoreactive axon terminals in other brain regions, further to those already indicated in Fig 6 (n=3). For each image, distance from bregma (in mm) is indicated at the bottom right. Scale bars = 100 µm. 40x magnification. Abbreviations: AHA: anterior hypothalamic area; ARC: arcuate nucleus; DMH: dorsomedial hypothalamus; EAC: extended amygdala central part; LC: locus coeruleus; LHA: lateral hypothalamic area; LPBN: lateral parabrachial nucleus; MPBN: medial parabrachial nucleus; PMNd: premammillary nucleus dorsal part; SI: substantia innominata; SuM: supramammillary nucleus; VTA: ventral tegmental area.

**Suppl. Fig 4:**
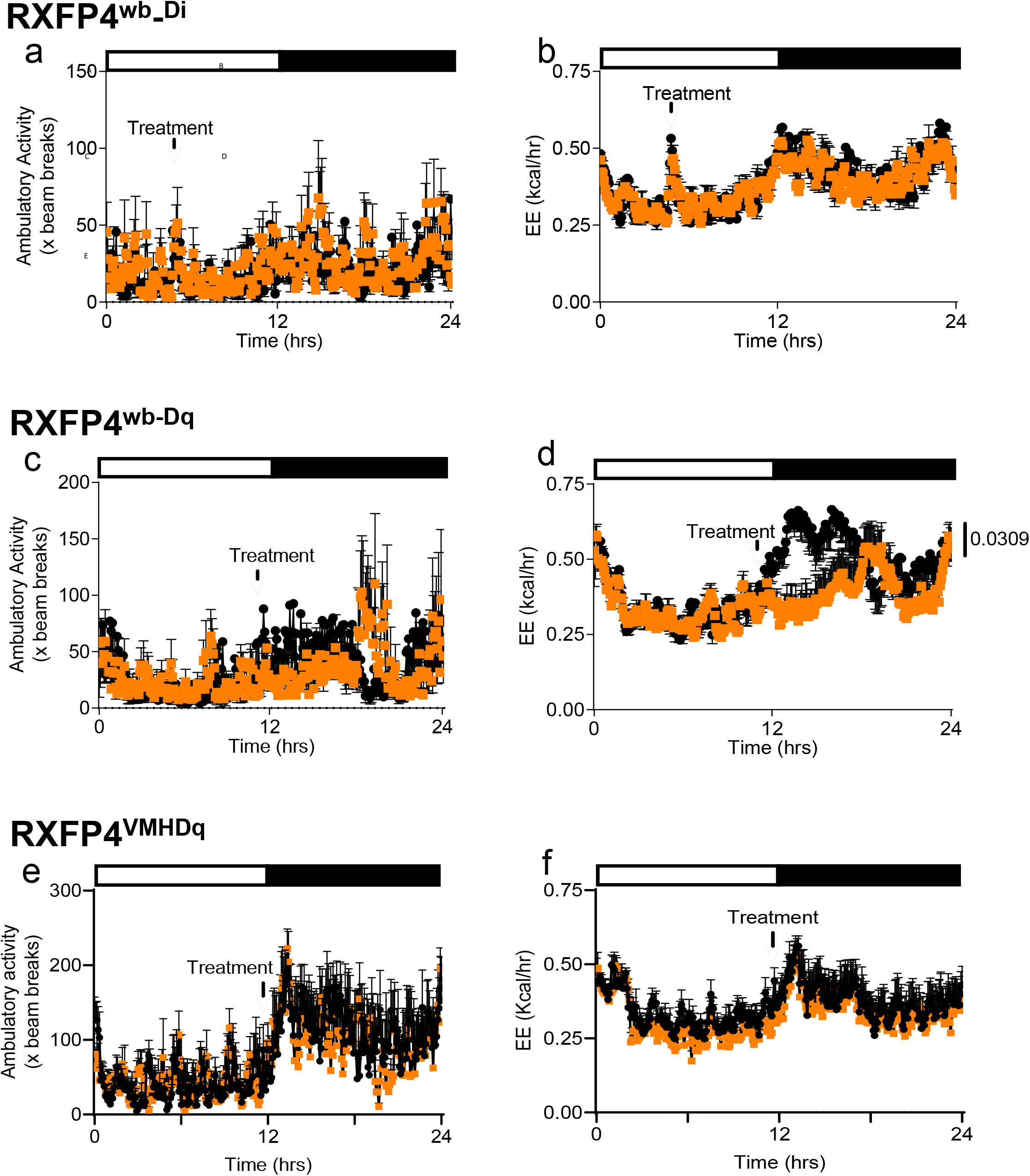
Calorimetric data from RXFP4 Di and Dq mice. Ambulatory activity (a,c,e) and energy expenditure (b,d,f) of RXFP4^wb-Di^ (a,b, effect of time [F(8.262, 115.7) = 2.092, p = 0.0402] and effect of time [F(11.52, 161.3) = 8.018, p < 0.0001] respectively), RXFP4^wb-Dq^ (c,d, effect of time [F(3.934,55.08) = 1.610, p = 0.0137 and effect of treatment [F(1, 13) = 5.860, p = 0.0309) and RXFP4^VMHDq^ (e,f, effect of time [F(288, 4018) = 3.622, p < 0.0001) and effect of time [F(4.194, 58.01) = 4.400, p = 0.0031) mice housed in metabolic cages in response to CNO (orange) or saline (black). Conditions and stats as in Fig 2 (Di), Fig 3 (Dq) and Fig 5 (VMH-Dq), respectively.

## Notes

### Summary of Updates

We have added extra data showing that Insl5 has direct effects on Rxfp4-Cre labeled cells in the VMH in acute slices and that intra-parenchymal application of Insl5 in the VMH affects food intake. We also ablated Rxfp4 neurons in the VMH to assess chronic outcomes. The manuscript has been rewritten/focused and in this context data concerning peripheral Rxfp4 expression has been removed and will be part of a different manuscript.

## References

1. Bathgate RA, Halls ML, van der Westhuizen ET, Callander GE, Kocan M, Summers RJ. Relaxin family peptides and their receptors. Physiol Rev. 2013;93(1):405–80.

2. Ang SY, Hutchinson DS, Evans BA, Hossain MA, Patil N, Bathgate RA, et al. The actions of relaxin family peptides on signal transduction pathways activated by the relaxin family peptide receptor RXFP4. Naunyn Schmiedebergs Arch Pharmacol. 2017;390(1):105–11.

3. Liu C, Lovenberg TW. Relaxin-3, INSL5, and their receptors. Results Probl Cell Differ. 2008;46:213–37.

4. Grosse J, Heffron H, Burling K, Akhter Hossain M, Habib AM, Rogers GJ, et al. Insulin-like peptide 5 is an orexigenic gastrointestinal hormone. Proceedings of the National Academy of Sciences of the United States of America. 2014;111(30):11133–8.

5. Lewis JE, Miedzybrodzka EL, Foreman RE, Woodward ORM, Kay RG, Goldspink DA, et al. Selective stimulation of colonic L cells improves metabolic outcomes in mice. Diabetologia. 2020;63(7):1396–407.

6. Billing LJ, Smith CA, Larraufie P, Goldspink DA, Galvin S, Kay RG, et al. Co-storage and release of insulin-like peptide-5, glucagon-like peptide-1 and peptideYY from murine and human colonic enteroendocrine cells. Mol Metab. 2018.

7. Lee YS, De Vadder F, Tremaroli V, Wichmann A, Mithieux G, Bäckhed F. Insulin-like peptide 5 is a microbially regulated peptide that promotes hepatic glucose production. Mol Metab. 2016;5(4):263–70.

8. Zaykov AN, Gelfanov VM, Perez-Tilve D, Finan B, DiMarchi RD. Insulin-like peptide 5 fails to improve metabolism or body weight in obese mice. Peptides. 2019;120:170116.

9. Liu C, Kuei C, Sutton S, Chen J, Bonaventure P, Wu J, et al. INSL5 is a high affinity specific agonist for GPCR142 (GPR100). J Biol Chem. 2005;280(1):292–300.

10. Boels K, Schaller HC. Identification and characterisation of GPR100 as a novel human G-protein-coupled bradykinin receptor. Br J Pharmacol. 2003;140(5):932–8.

11. Liu C, Chen J, Sutton S, Roland B, Kuei C, Farmer N, et al. Identification of relaxin-3/INSL7 as a ligand for GPCR142. J Biol Chem. 2003;278(50):50765–70.

12. Musatov S, Chen W, Pfaff DW, Mobbs CV, Yang XJ, Clegg DJ, et al. Silencing of estrogen receptor alpha in the ventromedial nucleus of hypothalamus leads to metabolic syndrome. Proc Natl Acad Sci U S A. 2007;104(7):2501–6.

13. Luo SX, Huang J, Li Q, Mohammad H, Lee CY, Krishna K, et al. Regulation of feeding by somatostatin neurons in the tuberal nucleus. Science. 2018;361(6397):76–81.

14. Muntean BS, Zucca S, MacMullen CM, Dao MT, Johnston C, Iwamoto H, et al. Interrogating the Spatiotemporal Landscape of Neuromodulatory GPCR Signaling by Real-Time Imaging of cAMP in Intact Neurons and Circuits. Cell Rep. 2018;22(1):255–68.

15. Kim DW, Yao Z, Graybuck LT, Kim TK, Nguyen TN, Smith KA, et al. Multimodal Analysis of Cell Types in a Hypothalamic Node Controlling Social Behavior. Cell. 2019;179(3):713–28.e17.

16. van Veen JE, Kammel LG, Bunda PC, Shum M, Reid MS, Massa MG, et al. Hypothalamic estrogen receptor alpha establishes a sexually dimorphic regulatory node of energy expenditure. Nat Metab. 2020;2(4):351–63.

17. Li MM, Madara JC, Steger JS, Krashes MJ, Balthasar N, Campbell JN, et al. The Paraventricular Hypothalamus Regulates Satiety and Prevents Obesity via Two Genetically Distinct Circuits. Neuron. 2019;102(3):653–67.e6.

18. Wall NR, Wickersham IR, Cetin A, De La Parra M, Callaway EM. Monosynaptic circuit tracing in vivo through Cre-dependent targeting and complementation of modified rabies virus. Proc Natl Acad Sci U S A. 2010;107(50):21848–53.

19. Jendryka M, Palchaudhuri M, Ursu D, van der Veen B, Liss B, Kätzel D, et al. Pharmacokinetic and pharmacodynamic actions of clozapine-N-oxide, clozapine, and compound 21 in DREADD-based chemogenetics in mice. Sci Rep. 2019;9(1):4522.

20. Woodward ORM, Gribble FM, Reimann F, Lewis JE. Gut peptide regulation of food intake – evidence for the modulation of hedonic feeding. The Journal of physiology. 2021.

21. Mashima H, Ohno H, Yamada Y, Sakai T, Ohnishi H. INSL5 may be a unique marker of colorectal endocrine cells and neuroendocrine tumors. Biochem Biophys Res Commun. 2013;432(4):586–92.

22. Thanasupawat T, Hammje K, Adham I, Ghia JE, Del Bigio MR, Krcek J, et al. INSL5 is a novel marker for human enteroendocrine cells of the large intestine and neuroendocrine tumours. Oncol Rep. 2013;29(1):149–54.

23. Ebling FJP, Lewis JE. Tanycytes and hypothalamic control of energy metabolism. Glia. 2018;66(6):1176–84.

24. Rogge G, Jones D, Hubert GW, Lin Y, Kuhar MJ. CART peptides: regulators of body weight, reward and other functions. Nat Rev Neurosci. 2008;9(10):747–58.

25. Cavuoto P, Wittert GA. The role of the endocannabinoid system in the regulation of energy expenditure. Best Pract Res Clin Endocrinol Metab. 2009;23(1):79–86.

26. Xu B, Goulding EH, Zang K, Cepoi D, Cone RD, Jones KR, et al. Brain-derived neurotrophic factor regulates energy balance downstream of melanocortin-4 receptor. Nat Neurosci. 2003;6(7):736–42.

27. Sampson CM, Kasper JM, Felsing DE, Raval SR, Ye N, Wang P, et al. Small-Molecule Neuromedin U Receptor 2 Agonists Suppress Food Intake and Decrease Visceral Fat in Animal Models. Pharmacol Res Perspect. 2018;6(5):e00425.

28. Moon YS, Smas CM, Lee K, Villena JA, Kim KH, Yun EJ, et al. Mice lacking paternally expressed Pref-1/Dlk1 display growth retardation and accelerated adiposity. Mol Cell Biol. 2002;22(15):5585–92.

29. Wermter AK, Scherag A, Meyre D, Reichwald K, Durand E, Nguyen TT, et al. Preferential reciprocal transfer of paternal/maternal DLK1 alleles to obese children: first evidence of polar overdominance in humans. Eur J Hum Genet. 2008;16(9):1126–34.

30. Viskaitis P, Irvine EE, Smith MA, Choudhury AI, Alvarez-Curto E, Glegola JA, et al. Modulation of SF1 Neuron Activity Coordinately Regulates Both Feeding Behavior and Associated Emotional States. Cell Rep. 2017;21(12):3559–72.

31. Wang D, He X, Zhao Z, Feng Q, Lin R, Sun Y, et al. Whole-brain mapping of the direct inputs and axonal projections of POMC and AgRP neurons. Front Neuroanat. 2015;9:40.

32. Williams G, Bing C, Cai XJ, Harrold JA, King PJ, Liu XH. The hypothalamus and the control of energy homeostasis: different circuits, different purposes. Physiology & behavior. 2001;74(4-5):683–701.

33. Kelley AE, Baldo BA, Pratt WE, Will MJ. Corticostriatal-hypothalamic circuitry and food motivation: integration of energy, action and reward. Physiology & behavior. 2005;86(5):773–95.

34. Kenny PJ. Reward mechanisms in obesity: new insights and future directions. Neuron. 2011;69(4):664–79.

35. Tobiansky DJ, Roma PG, Hattori T, Will RG, Nutsch VL, Dominguez JM. The medial preoptic area modulates cocaine-induced activity in female rats. Behav Neurosci. 2013;127(2):293–302.

36. Tryon VL, Mizumori SJY. A Novel Role for the Periaqueductal Gray in Consummatory Behavior. Front Behav Neurosci. 2018;12:178.

37. Wu Q, Boyle MP, Palmiter RD. Loss of GABAergic signaling by AgRP neurons to the parabrachial nucleus leads to starvation. Cell. 2009;137(7):1225–34.

38. Chiang MC, Bowen A, Schier LA, Tupone D, Uddin O, Heinricher MM. Parabrachial Complex: A Hub for Pain and Aversion. J Neurosci. 2019;39(42):8225–30.

39. Lo L, Yao S, Kim DW, Cetin A, Harris J, Zeng H, et al. Connectional architecture of a mouse hypothalamic circuit node controlling social behavior. Proc Natl Acad Sci U S A. 2019;116(15):7503–12.

40. King BM. The rise, fall, and resurrection of the ventromedial hypothalamus in the regulation of feeding behavior and body weight. Physiol Behav. 2006;87(2):221–44.

41. Miller NE, Bailey CJ, Stevenson JA. Decreased “hunger” but increased food intake resulting from hypothalamic lesions. Science. 1950;112(2905):256–9.

42. Teitelbaum P. Random and food-directed activity in hyperphagic and normal rats. J Comp Physiol Psychol. 1957;50(5):486–90.

43. Grossman SP. The VMH: a center for affective reactions, satiety, or both? . 1966;1:1–10.

44. Lindberg D, Chen P, Li C. Conditional viral tracing reveals that steroidogenic factor 1-positive neurons of the dorsomedial subdivision of the ventromedial hypothalamus project to autonomic centers of the hypothalamus and hindbrain. J Comp Neurol. 2013;521(14):3167–90.

45. Hashikawa K, Hashikawa Y, Tremblay R, Zhang J, Feng JE, Sabol A, et al. Esr1. Nat Neurosci. 2017;20(11):1580–90.

46. Coutinho EA, Okamoto S, Ishikawa AW, Yokota S, Wada N, Hirabayashi T, et al. Activation of SF1 Neurons in the Ventromedial Hypothalamus by DREADD Technology Increases Insulin Sensitivity in Peripheral Tissues. Diabetes. 2017;66(9):2372–86.

47. Yang CF, Chiang MC, Gray DC, Prabhakaran M, Alvarado M, Juntti SA, et al. Sexually dimorphic neurons in the ventromedial hypothalamus govern mating in both sexes and aggression in males. Cell. 2013;153(4):896–909.

48. Lee H, Kim DW, Remedios R, Anthony TE, Chang A, Madisen L, et al. Scalable control of mounting and attack by Esr1+ neurons in the ventromedial hypothalamus. Nature. 2014;509(7502):627–32.

49. Krause WC, Ingraham HA. Origins and Functions of the Ventrolateral VMH: A Complex Neuronal Cluster Orchestrating Sex Differences in Metabolism and Behavior. Adv Exp Med Biol. 2017;1043:199–213.

50. Correa SM, Newstrom DW, Warne JP, Flandin P, Cheung CC, Lin-Moore AT, et al. An estrogen-responsive module in the ventromedial hypothalamus selectively drives sex-specific activity in females. Cell Rep. 2015;10(1):62–74.

51. Ma S, Smith CM, Blasiak A, Gundlach AL. Distribution, physiology and pharmacology of relaxin-3/RXFP3 systems in brain. Br J Pharmacol. 2017;174(10):1034–48.

52. Atasoy D, Aponte Y, Su HH, Sternson SM. A FLEX switch targets Channelrhodopsin-2 to multiple cell types for imaging and long-range circuit mapping. J Neurosci. 2008;28(28):7025–30.

53. Shimshek DR, Kim J, Hübner MR, Spergel DJ, Buchholz F, Casanova E, et al. Codon-improved Cre recombinase (iCre) expression in the mouse. Genesis. 2002;32(1):19–26.

54. Srinivas S, Watanabe T, Lin CS, William CM, Tanabe Y, Jessell TM, et al. Cre reporter strains produced by targeted insertion of EYFP and ECFP into the ROSA26 locus. BMC Dev Biol. 2001;1:4.

55. Zhu H, Aryal DK, Olsen RH, Urban DJ, Swearingen A, Forbes S, et al. Cre-dependent DREADD (Designer Receptors Exclusively Activated by Designer Drugs) mice. Genesis. 2016;54(8):439–46.

56. Zariwala HA, Borghuis BG, Hoogland TM, Madisen L, Tian L, De Zeeuw CI, et al. A Cre-dependent GCaMP3 reporter mouse for neuronal imaging in vivo. J Neurosci. 2012;32(9):3131–41.

57. Luche H, Weber O, Nageswara Rao T, Blum C, Fehling HJ. Faithful activation of an extra-bright red fluorescent protein in “knock-in” Cre-reporter mice ideally suited for lineage tracing studies. Eur J Immunol. 2007;37(1):43–53.

58. Heath CJ, Phillips BU, Bussey TJ, Saksida LM. Measuring Motivation and Reward-Related Decision Making in the Rodent Operant Touchscreen System. Current protocols in neuroscience. 2016;74:8.34.1–8..20.

59. Adriaenssens AE, Biggs EK, Darwish T, Tadross J, Sukthankar T, Girish M, et al. Glucose-Dependent Insulinotropic Polypeptide Receptor-Expressing Cells in the Hypothalamus Regulate Food Intake. Cell metabolism. 2019;30(5):987–96.e6.

60. Picelli S, Björklund Å, Faridani OR, Sagasser S, Winberg G, Sandberg R. Smart-seq2 for sensitive full-length transcriptome profiling in single cells. Nat Methods. 2013;10(11):1096–8.

