## Supplementary material for "Relaxin/insulin-like family peptide receptor 4 (Rxfp4) expressing hypothalamic neurons modulate food intake and preference in mice": suppl methods

**Suppl. Methods 1: *Rxfp4-Cre mouse generation***

To express Cre recombinase under the control of the *Rxfp4* promoter, we replaced the sequence between the start codon and the stop codon in the single exon coding for *Rxfp4* in the murine-based BAC RP24-72I4 (Children’s Hospital Oakland Research Institute) initially by a counter-selection cassette rpsL-neo (GeneBridges) and subsequently by the improved Cre (iCre) sequence using Red/ET recombination technology (GeneBridges) (see Fig. 1a). Briefly, the rpsL-neo {*Rxfp4-001/2*} or iCre {*Rxfp4-005/6*} sequences were amplified by PCR with Phusion-polymerase with the the primers given in {}, adding Rxfp4-gene specific 3′ and 5′ sequences; the sequences of the primers are given in Suppl. Method Table 1 below. Homologous recombination was achieved upon co-transforming the BAC containing Escherichia coli DH10B clone with the PCR product and the plasmid pSC101-BAD-gbaA, which provides the recombination enzymes (GeneBridges). Positive recombinants were isolated using appropriate antibiotic selection and characterized by PCR (primer pairs: *Rxfp4-003/iCre-003* (796 bp); *Rxfp4-004/iCre-002* (1115 bp)) and restriction analysis (EagI; AatII and XhoI-digest fingerprints). Identity and correct positioning of the introduced iCre sequence was confirmed by direct sequencing using the oligonucleotides Rxfp4-003, Rxfp4-004, iCre001 and iCre003. BAC DNA for microinjection was purified using the large-construct Maxi-Prep kit (Qiagen) and dissolved at ∼1–2 ng/μL in injection buffer containing (mmol/L): 10 Tris-HCl (pH 7.5), 0.1 EDTA, 100 NaCl, 0.03 spermine, and 0.07 spermidine. Pronuclear injection into ova derived from C57B6/CBA F1 parents and reimplantation of embryos into pseudopregnant females was performed by the Central Biomedical Services at Cambridge University. DNA of pups was isolated from ear clips by proteinase K digestion and screened for the transgene by PCR using the following primer pairs: *Rxfp4-003/iCre003*, *Rxfp4-004/iCre002,* *iCre002/003* and *RM41/42 (β-catenin )*, with the latter serving as a DNA quality control. Initially we received four potential founders, however, only two of these passed the transgene on to their offsprings (“Rxfp4-70” and “Rxfp4-73”). Both showed detectable reporter expression when crossed with a Rosa26-fxSTOPfx-reporter lines. The transgene copy-number for the Rxfp4-73 strain was estimated to be n=1 by quantitative PCR using iCre and Kcnj11-primer/probes. The Rxfp4-73 founder strain was backcrossed for more than eight generations onto a C57B6 background.

**A**

Suppl. Methods Table 1: Oligonucleotides used to make/verify Rxfp4-Cre BAC

| Name | Sequence |
| --- | --- |
| Rxfp4-001  (-rpsLneo) | AGG CGG GCA CTC CCT GGT TCC TCT GCT CTG CTG TGC TCT AGC AAC CTC CGC GGT CTT GCG ATG GGC CTG GTG ATG ATG GCG GGA TCG |
| Rxfp4-002  (-rpsLneo) | CAC CTC CTG TTT GGA CTC CAG CTT CTC CGT TCT GTC ACA CCC AGA GTT GGT GAT CAA AGA TCA GAA GAA CTC GTC AAG AAG GCG |
| Rxfp4-003 | CCA ACA CCC AGA TTC CAA G (e.g. to screen recombinant BACs with rpsLneo002; exp: 1459 bp) |
| Rxfp4-004 | CGG GTG ATG GAC GTT AGT TAT C (e.g. to screen recombinant BACs with rpsLneo001; exp: 1499 bp) |
| RpsLneo001 | CCT GGT GAT GAT GGC G |
| RpsLneo002 | AAG AAC TCG TCA AGA AGG CG |
| Rxfp4-005  (-iCRE) | AGG CGG GCA CTC CCT GGT TCC TCT GCT CTG CTG TGC TCT AGC AAC CTC CGC GGT CTT GCG ATG GTG CCC AAG AAG AAG AGG |
| Rxfp4-006  (-iCRE) | CAC CTC CTG TTT GGA CTC CAG CTT CTC CGT TCT GTC ACA CCC AGA GTT GGT GAT CAA AGA TCA GTC CCC ATC CTC GAG CAG C |
| iCre001 | GTA GTC CCT CAC ATC CTC AGG |
| iCre002 | GAC AGG CAG GCC TTC TCT GAA |
| iCre003 | CTT CTC CAC ACC AGC TGT GGA |
| iCre004-fw | GCC GAA ATT GCC AGA ATC AG |
| iCre005-rev | CAA TGT GGA TCA GCA TTC TCC |
| iCre-probe | 6-fam-TGA AGG ACA TCT CCC GCA CCG-tAMRA |
| RM41 | AAG GTA GAG TGA TGA AAG TTG TT |
| RM41 | CAC CAT GTC CTC TGT CTA TTC |
| mKcnj11-fw | ccc gct tcg tgt cca aga |
| mKcnj11-rev | cag cgt ggt gaa cac atc ct |
| mKcnj11-probe | 6-fam-caa cgt cgc cca caa gaa cat tcg a-BHQ-1 |
