## Supplementary figures and images for "Relaxin/insulin-like family peptide receptor 4 (Rxfp4) expressing hypothalamic neurons modulate food intake and preference in mice"

### suppl video

#### Slide 1
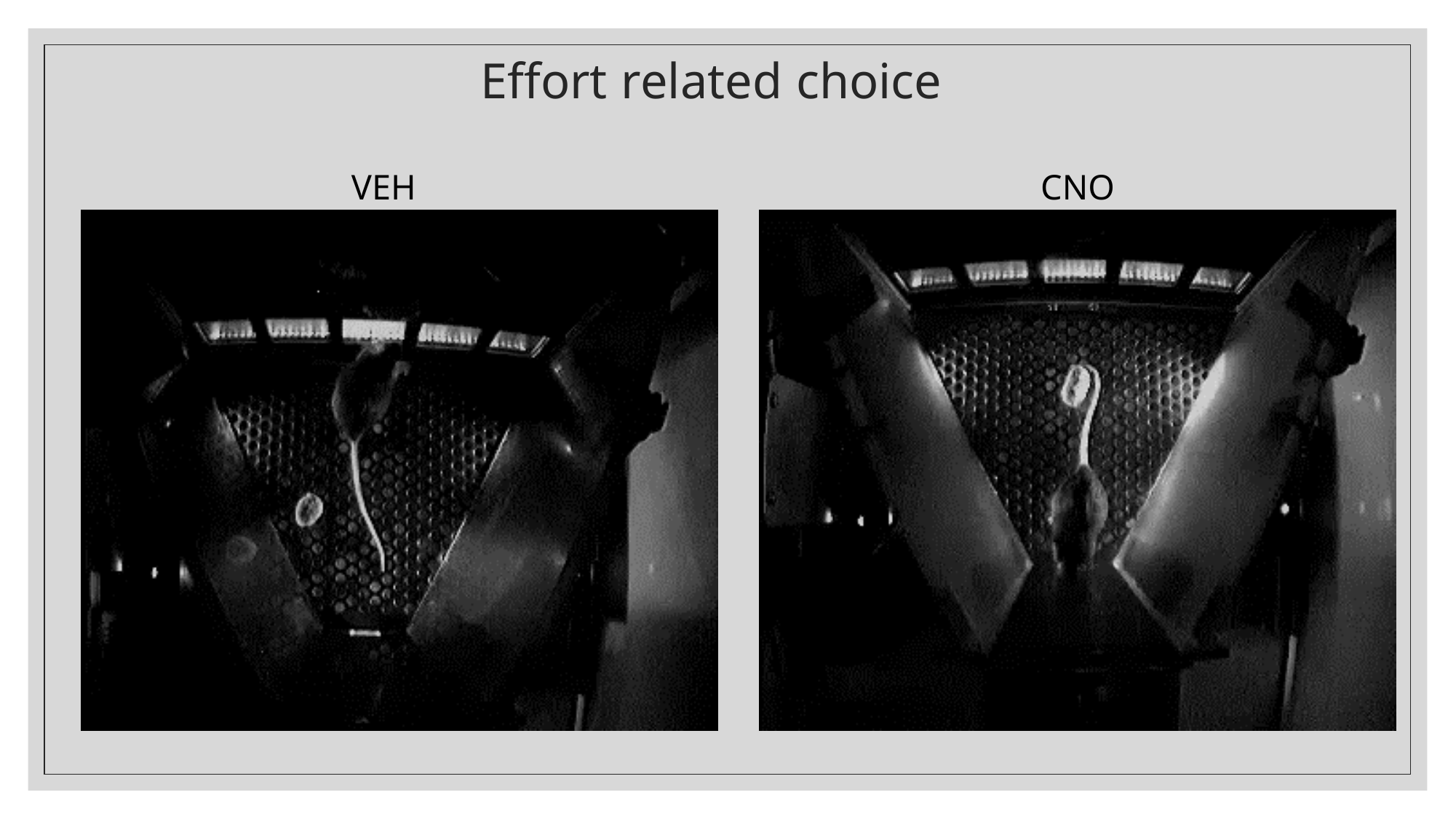

##### Effort related choice
VEH
CNO
